# Unconventional secretion of unglycosylated ORF8 is critical for the cytokine storm during SARS-CoV-2 infection

**DOI:** 10.1101/2021.12.03.471057

**Authors:** Xiaoyuan Lin, Beibei Fu, Yan Xiong, Na Xing, Weiwei Xue, Dong Guo, Mohamed Y. Zaky, Krishna Chaitanya Pavani, Dusan Kunec, Jakob Trimpert, Haibo Wu

## Abstract

Coronavirus disease 2019 is a respiratory infectious disease caused by the severe acute respiratory syndrome coronavirus 2 (SARS-CoV-2). Evidence on the pathogenesis of SARS-CoV-2 is accumulating rapidly. In addition to structural proteins such as Spike and Envelope, the functional roles of non-structural and accessory proteins in regulating viral life cycle and host immune responses remain to be understood. Here, we show that open reading frame 8 (ORF8) acts as messenger for inter-cellular communication between alveolar epithelial cells and macrophages during SARS-CoV-2 infection. Mechanistically, ORF8 is a secretory protein that can be secreted by infected epithelial cells via both conventional and unconventional secretory pathways. The unconventionally secreted ORF8 recognizes the IL17RA receptor of macrophages and induces cytokine release. However, conventionally secreted ORF8 cannot bind to IL17RA due to N-linked glycosylation. Furthermore, we found that Yip1 interacting factor homolog B (YIF1B) is a channel protein that translocates unglycosylated ORF8 into vesicles for unconventional secretion. Blocking the unconventional secretion of ORF8 via a YIF1B knockout in hACE2 mice attenuates inflammation and yields delayed mortality following SARS-CoV-2 challenge.

## Introduction

Coronavirus disease 2019 (COVID-19), caused by severe acute respiratory syndrome coronavirus 2 (SARS-CoV-2), is continuing to spread around the world with nearly 256 million confirmed cases and more than 5.1 million deaths. According to clinical case reports, critically ill patients with COVID-19 experience a cytokine storm, resulting in acute respiratory distress syndrome and multiple organ failure(*1, 2*). Currently, our understanding of the mechanisms behind the cytokine storm and how SARS-CoV-2 affects cytokine release is still limited.

Open reading frame 8 (ORF8) is an accessory protein of SARS-CoV-2 and it is one of the most rapidly evolving β-coronaviruses proteins(*3*). A 29 nucleotide deletion in ORF8 is the most obvious genetic change in severe acute respiratory syndrome coronavirus (SARS or SARS-CoV-1) during its host-jump from bats to humans(*4*). The Δ382 variant of SARS-CoV-2, which eliminates ORF8 transcription, seems to be associated with milder infection and less systemic release of pro-inflammatory cytokines(*5*). In a previous study, we demonstrated that ORF8 contributes to the cytokine storm during SARS-CoV-2 infection(*6*). Specifically, we found ORF8 to interact with the IL17RA receptor, leading to excessive activation of IL-17 signaling and downstream NF-κB pathway. However, it remains unclear how the virus exposes ORF8 to enable access to the extracellular domain of IL17RA.

In eukaryotes, secretory proteins usually contain a signal peptide that triggers translocation into the endoplasmic reticulum (ER)(*7*). Following this translocation to the ER, cargoes will be exported through ER-Golgi trafficking for further processing and modification(*8, 9*). This process is termed conventional secretion. Besides, many cytosolic proteins without signal peptides, such as fibroblast growth factor 2 and yeast Acb1, can be released through an unconventional protein secretion pathway(*10, 11*). Recently, Zhang et al. identified TMED10 as a protein channel for vesicle entry and secretion of interleukin 1 family members(*12*). Evidently, viral proteins can hijack this secretion pathway to become secreted. For example, the HIV-1 Nef protein can be released from infected cells via an exosomal pathway(*13, 14*). However, very little is known about whether and how SARS-CoV-2 encoded proteins are secreted during infection.

Glycosylation is a common posttranslational modification, it involves the addition of glycans to macromolecules and is considered essential for the correct folding and functional performance of proteins(*15–17*). It is also not uncommon for proteins from pathogens to be glycosylated by the host. The co-evolution of N-linked glycosylation sites in influenza viruses affects the host specificity(*18*). The glycosylation of viral envelope proteins has a wide range of functions, including regulating cell tropism, protein stability and immune evasion(*19–21*). Recent studies have shown that SARS-CoV-2 Spike protein has 22 N-linked glycosylation sites and 17 O-linked glycosylation sites, which may influence viral infectivity and pathogenicity(*22–24*). In contrast to the situation for the Spike protein, glycosylation and its functional role in accessory proteins of SARS-CoV-2 has not yet been reported.

Here, we identified SARS-CoV-2 ORF8 as a secretory protein that can be secreted via conventional and unconventional secretory pathways at the same time. We found that unglycosylated ORF8 secreted via an unconventional pathway is responsible for the release of pro-inflammatory cytokines by binding the IL17RA receptor. By contrast, conventionally secreted ORF8 is incapable of binding to IL17RA due to the N-linked glycosylation at Asn78 site. Further, we identified Yip1 interacting factor homolog B (YIF1B) as a channel protein that recognizes and translocates unglycosylated ORF8 into vesicles, thus enabling unconventional secretion. Our findings present an important contribution to the understanding of how SARS-CoV-2 promotes the onset of cytokine storm, and provide a promising strategy for the development of COVID-19 therapeutics.

## Results

### SARS-CoV-2 ORF8 is a secretory protein that is associated with cytokine release

In order to investigate the secretion of ORF8 protein, we infected Calu-3 human lung epithelial cells with SARS-CoV-2 that was generated using a reverse genetic system (*25–27*)(Fig. 1A). Cell culture supernatant was collected and presence of ORF8 was determined by ELISA and western blotting (Fig. 1B). We found that ORF8 protein can be secreted into cell culture medium (Fig. 1C). To further validate these results, we used the non-secretory SARS-CoV-2 main proteinase (M pro, also known as 3CL pro) non-secreted control, and the well-known Nef secretory protein of HIV-1 (*13, 14*) as a secreted protein control. We found that Jurkat cells secreted Nef protein following HIV-1 infection, and Calu-3 cells secreted ORF8 protein after being infected with SARS-CoV-2 (Fig. 1D). By contrast, the structural protein 3CL pro was not secreted (Fig. 1D). Next, we tested the time-dependency of ORF8 secretion in Calu-3 epithelial cells. After 12 hours of SARS-CoV-2 infection, the cell culture medium was replaced and ORF8 protein in the supernatant was detected every 2 hours. Using 3CL pro as a negative control, we found that the secretion of ORF8 continued for at least 12 hours after replacing the culture medium (Fig. 1E). These results indicated that SARS-CoV-2 ORF8 is a secretory protein.

**Figure 1.**
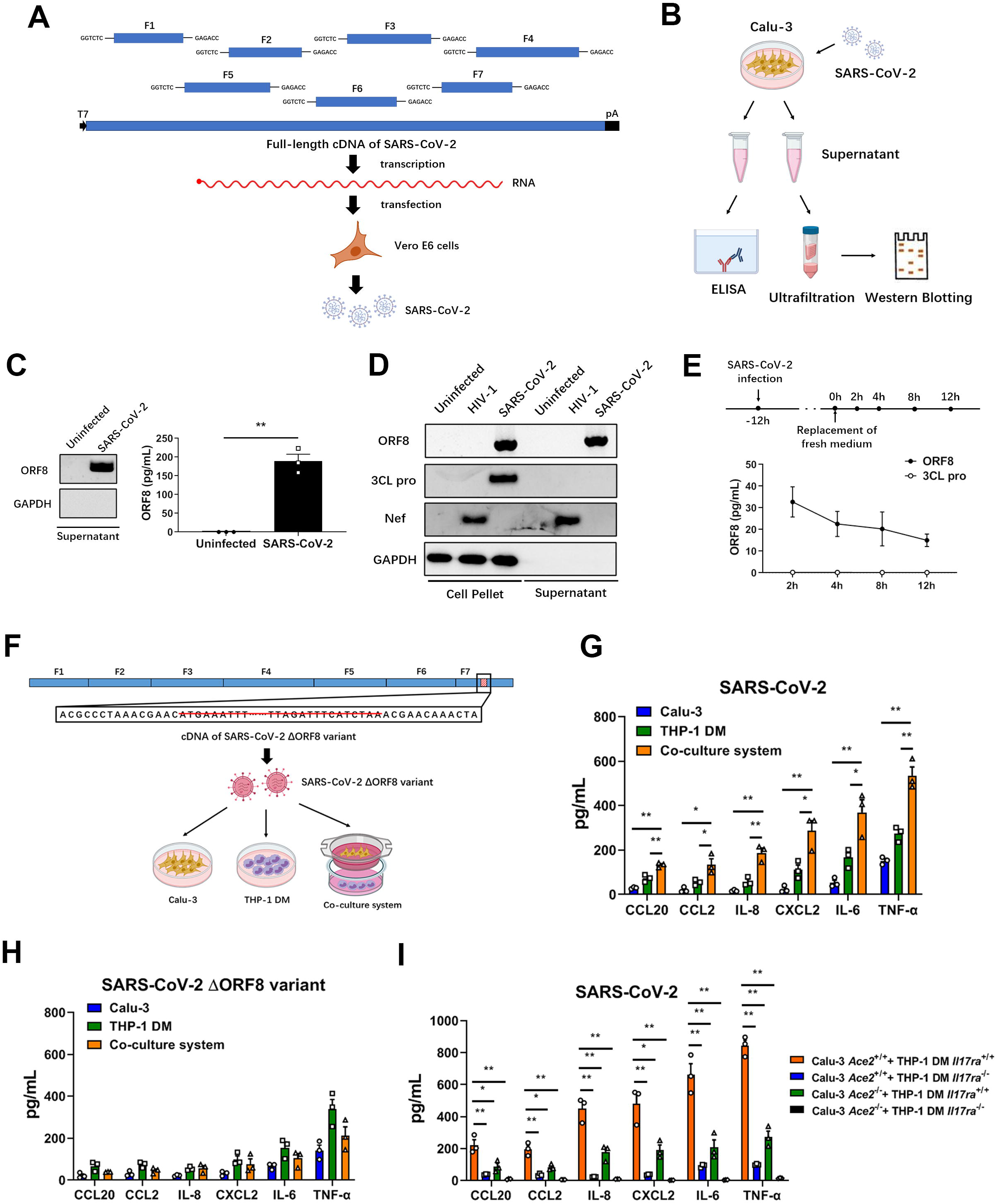
SARS-CoV-2 ORF8 can be secreted by epithelial cells. (A) Schematic diagram of SARS-CoV-2 infectious cDNA clone generated by a reverse genetic system. The cDNA fragments F1-F7 were synthesized and assembled into full-length SARS-CoV-2 cDNA, and RNA transcription, electroporation, and virus production were carried out in Vero E6 cells. (B) Schematic diagram of SARS-CoV-2 infection model. Calu-3 epithelial cells were infected with SARS-CoV-2 for 12 hours at a dosage of 10^5^ TCID_50_/mL. Cell culture supernatant was centrifuged and divided into two parts for western blotting and ELISA, respectively. (C) The secretion of ORF8 in (B) was detected by western blotting and ELISA. Representative images from n = 3 biological replicates are shown. Data in histogram are shown as the mean ± s.e.m. of n = 3 biological replicates. (D) Jurkat cells and Calu-3 epithelial cells were infected with HIV-1 and SARS-CoV-2 for 12 hours at a dosage of 10^5^ TCID_50_/mL. The secretion of ORF8, Nef and 3CL pro was detected by western blotting. Representative images from n = 3 biological replicates are shown. (E) Schematic diagram of time-dependent ORF8 secretion upon SARS-CoV-2 infection. After 12 hours of SARS-CoV-2 infection, cell culture medium was replaced, and the amount of ORF8 protein in the supernatant was detected by ELISA every 2 hours. Data are shown as the mean ± s.e.m. of n = 3 biological replicates. (F-H) Schematic diagram of ORF8-deleted SARS-CoV-2 variant (F). ORF8 coding sequence was deleted from the cDNA of F7 fragment. SARS-CoV-2 ΔORF8 variant was used to infect Calu-3 cells, THP-1 DM cells, and the co-culture system at a dosage of 10^5^ TCID_50_/mL. The secretion of cytokines and chemokines related to cytokine storm was detected by ELISA (G, H). Data are shown as the mean ± s.e.m. of n = 3 biological replicates (G, H). (H) Calu-3 *Ace2*^+/+^, Calu-3 *Ace2*^-/-^, THP-1 DM *Il17ra*^+/+^ and THP-1 DM *Il17ra*^-/-^ cells were used to form four kinds of culture systems. The secretion of cytokines and chemokines in different cell culture systems was detected by ELISA. Data are shown as the mean ± s.e.m. of n = 3 biological replicates. Unpaired two-tailed Student *t* test (C) and one-way ANOVA followed by Bonferroni post *hoc* test (G, I) were used for data analysis. *, p < 0.05, **, p < 0.01.

Our previous study has shown that ORF8 protein contributes to the cytokine storm during SARS-CoV-2 infection(*6*). SARS-CoV-2 mainly invades alveolar epithelial cells through binding to ACE2 receptors, however monocytes/macrophages play a critical role in the secretion and regulation of cytokines. Next, we generated a SARS-CoV-2 variant with an ORF8 deletion (Fig. 1F), and constructed an epithelial cell-macrophage co-culture system using a Transwell setup (Fig. 1F), to answer the question that whether ORF8 is a key factor in modulating the transmission process of infection signals from epithelial cells to monocytes/macrophages. In this system, the release of pro-inflammatory factors in macrophages infected with either wild-type SARS-CoV-2 or ORF8-deletion variant was increased, probably due to a small amount of ACE2 receptor expressed on the surface of macrophages (Fig. 1G, H). Interestingly, we found that the amount of pro-inflammatory factors in the co-culture system was much higher than that measured in individual epithelial cells or macrophages infected with wild-type SARS-CoV-2 (Fig. 1G). However, this synergistic pro-inflammatory effect between epithelial cells and macrophages was not observed in the ORF8-deletion virus infected group (Fig. 1H). Considering that ACE2 is responsible for virus entry(*28, 29*), and IL17RA is the receptor of ORF8(*6*), we generated ACE2-deficient epithelial cells (Calu-3 *Ace2*^-/-^) based on Calu-3 cell line, and IL17RA-deficient macrophages [THP-1-derived macrophages (THP-1 DM) *Il17ra*^-/-^] based on THP-1 cell line (Fig. S1A). By using these two ways to disrupt cellular communication between epithelial cells and macrophages, preventing SARS-CoV-2 entry by ACE2 deletion in Calu-3 *Ace2*^-/-^ cells, or interrupting ORF8 reception by IL17RA deletion in THP-1 DM *Il17ra*^-/-^ cells, a significant downregulation of pro-inflammatory factors was observed in the co-culture system (Fig. 1H). These results implied that inter-cellular communication between epithelial cells and monocytes/macrophages is important for the cytokine release during SARS-CoV-2 infection.

It is known that secretory proteins, such as Nef, can be secreted in absence of viral infection in an *in vitro* system(*13*). We therefore tested whether ORF8 protein can be secreted by human embryonic kidney (HEK-293FT) cells transfected with an ORF8-Flag plasmid. The results showed that Flag-tagged ORF8 was secreted in absence of SARS-CoV-2 infection (Fig. S1B, C). Furthermore, the supernatant of cells transfected with ORF8-Flag was collected and added to the culture medium of THP-1 DM cells for stimulation. In this setup, the release of pro-inflammatory factors in THP-1 DM cells stimulated with ORF8-Flag transfection supernatant was significantly increased compared to cells treated with control supernatant (Fig. S1D). In summary, we demonstrated that SARS-CoV-2 ORF8 is a secretory protein, and that extracellular ORF8 protein is of greatly promoting cytokine release.

### SARS-CoV-2 ORF8 has an unconventional secretory pathway

In order to understand the secretion pattern of ORF8, we used an online tool, Simple Modular Architecture Research Tool (SMART, http://smart.embl-heidelberg.de), to analyze structural domains of the ORF8 protein. We found a hydrophobic central domain (similar to the conserved signal peptide of eukaryotes) to be located at its N-terminus. We defined this domain as the signal peptide of ORF8 protein. Signal peptide-deficient mutant (ΔSignal-SARS-CoV-2 ORF8) was constructed and transfected into Calu-3 epithelial cells and HEK-293FT cells, respectively (Fig. 2A). The result showed that the secretion ability of ORF8 was substantially impaired by signal peptide deletion (Fig. 2B, C). Interestingly, an appreciable quantity of signal peptide-deficient ORF8 was observed in both the supernatant of Calu-3 and HEK-293FT cells. This result strongly suggested that the secretion of ORF8 is not completely blocked by the signal peptide deletion (Fig. 2B, C). This data implied that secretion of ORF8 might not completely depend on the presence of the signal peptide.

**Figure 2.**
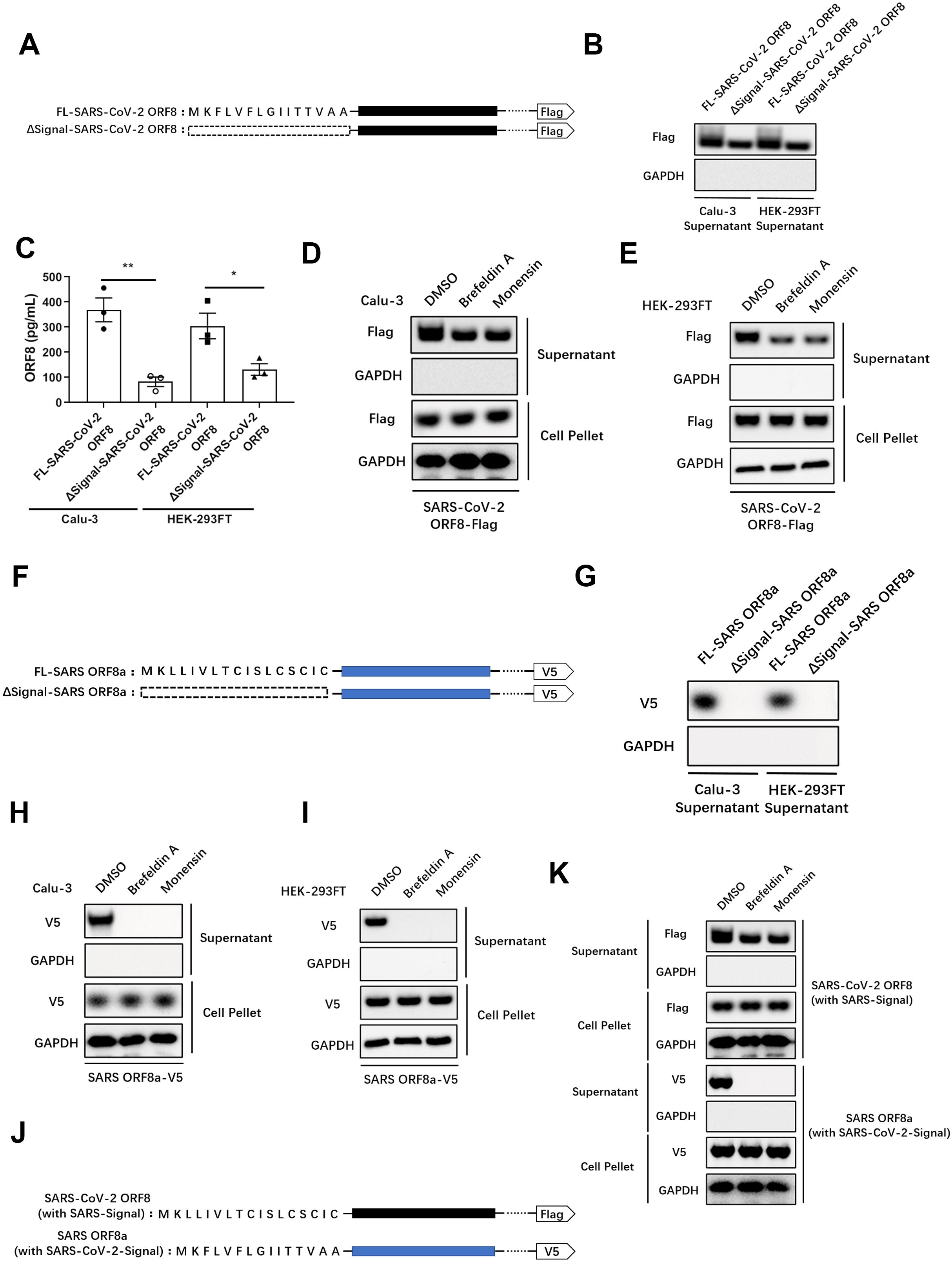
SARS-CoV-2 ORF8, rather than SARS ORF8, has an unconventional secretion pattern. (A) Structures of SARS-CoV-2 ORF8 and its signal peptide-deleted mutant. (B, C) Full-length SARS-CoV-2 ORF8 or its signal peptide-deleted mutant were transfected into Calu-3 cells or HEK-293FT cells. The secretion of ORF8 was detected by western blotting (B) and ELISA (C). Representative images from n = 3 biological replicates are shown (B). Data are shown as the mean ± s.e.m. of n = 3 biological replicates (C). (D, E) Brefeldin A (3 μg/mL) or Monensin (2 μM) was used to pretreat Calu-3 cells (D) or HEK-293FT cells (E) for 2 hours. Full-length SARS-CoV-2 ORF8 was transfected into pretreated cells. After 12 hours, the secretion of ORF8 was detected by western blotting. Representative images from n = 3 biological replicates are shown. (F) Structures of SARS ORF8a and its signal peptide-deleted mutant. (G) Full-length SARS ORF8a or its signal peptide-deleted mutant were transfected into Calu-3 cells or HEK-293FT cells. The secretion of ORF8a was detected by western blotting. Representative images from n = 3 biological replicates are shown. (H, I) Brefeldin A (3 μg/mL) or Monensin (2 μM) was used to pretreat Calu-3 cells (H) or HEK-293FT cells (I) for 2 hours. Full-length SARS ORF8a was transfected into pretreated cells. After 12 hours, the secretion of ORF8a was detected by western blotting. Representative images from n = 3 biological replicates are shown. (J) Structures of SARS-CoV-2 ORF8 mutant with signal peptide from SARS ORF8a (with SARS-Signal), and SARS ORF8a mutant with signal peptide from SARS-CoV-2 ORF8 (with SARS-CoV-2-Signal). (K) Brefeldin A (3 μg/mL) or Monensin (2 μM) was used to pretreat Calu-3 cells for 2 hours. SARS-CoV-2 ORF8 with SARS-Signal and SARS ORF8a with SARS-CoV-2-Signal were transfected into pretreated Calu-3 cells. After 12 hours, the secretion of ORF8 was detected by western blotting. Representative images from n = 3 biological replicates are shown. Unpaired two-tailed Student *t* test (C) was used for data analysis. *, p < 0.05, **, p < 0.01.

Proteins secreted through the conventional secretory pathway contain an N-terminal signal peptide, which is recognized by the signal recognition particles and transported into the ER, followed by signal peptide cleavage and trafficking to the Golgi apparatus and the subsequent endomembrane system(*7*). To verify the existence of an unconventional secretory pathway for ORF8, Brefeldin A(*30*) or Monensin(*31*) was used to inhibit the Golgi-related vesicle transportation. Consistent with our previous result, secretion of ORF8 was still observed when the conventional secretory pathway was blocked (Fig. 2D, E).

To understand whether the secretion pattern of ORF8 is evolutionary conserved, we analyzed the homologous ORF8 of SARS and found the SARS ORF8a isoform to contain an N-terminal signal peptide. We then constructed a ΔSignal-SARS ORF8a mutant and transfected it into epithelial cells (Fig. 2F). While the intact SARS ORF8a was secreted normally (Fig. 2G), the ΔSignal-SARS ORF8a lost the secretory ability completely (Fig. 2G). This finding stands in interesting contrast to our results showing that SARS-CoV-2 ORF8 is secreted in absence of the signal peptide (Fig. 2G). This finding was further supported by inhibition of the Golgi-related vesicle transport using Brefeldin A or Monensin in Calu-3 (Fig. 2H) and HEK-293FT cells (Fig. 2I). The results obtained here show that SARS ORF8a can only be secreted under the guidance of signal peptide through the conventional secretory pathway.

Considering that the signal peptides of these two different ORF8 proteins share distinct sequences, we asked whether the different secretion patterns of ORF8 is due to the different types of signal peptide. To examine this hypothesis, we exchanged the ORF8 signal peptides between SARS and SARS-CoV-2 (Fig. 2J). As a result, exchange of signal peptides did not change the secretory patterns observed for the two viruses (Fig. 2K). Specifically, SARS-CoV-2 ORF8 carrying the SARS ORF8a signal peptide was still secreted when the Golgi-dependent secretory pathway was blocked (Fig. 2K). Taken together, these results indicated that SARS-CoV-2 ORF8 is likely to have an unconventional secretion pattern that does not depend on the presence of a signal peptide.

### Unconventional secretion of ORF8 is required for cytokine storm

Next, we studied whether the different secretion patterns of ORF8 are associated with the release of pro-inflammatory factors. Calu-3 epithelial cells were pretreated with Brefeldin A or Monensin to block the conventional secretory pathway, then cells were infected with SARS-CoV-2 and culture supernatant was collected to stimulate THP-1 DM cells. After 12 hours, pro-inflammatory factors secreted by macrophages were examined by ELISA (Fig. 3A). Surprisingly, although the amount of secreted ORF8 was significantly decreased (Fig. 3B), the release of pro-inflammatory factors was barely affected by Brefeldin A or Monensin treatment (Fig. 3C). In order to rule out the possibility of mutual influence between macrophages themselves, we tested pro-inflammatory factors released by THP-1 DM cells stimulated with supernatant at different time intervals. The result showed that there were no significant differences in the secretion of pro-inflammatory factors by macrophages within 0-12 h (Fig. 3C, Fig. S2A). Further, we used exogenously expressed Flag-tagged ORF8 (Fig. 3A), instead of SARS-CoV-2 virus, to validate this finding. Consistent with previous results, we observed almost equal expression levels of pro-inflammatory factors induced by ORF8-Flag regardless of the Golgi-dependent secretory pathway blockade (Fig. 3D, Fig. S2B). These results implied that the unconventionally, instead of the conventionally secreted ORF8 is responsible for the cytokines release.

**Figure 3.**
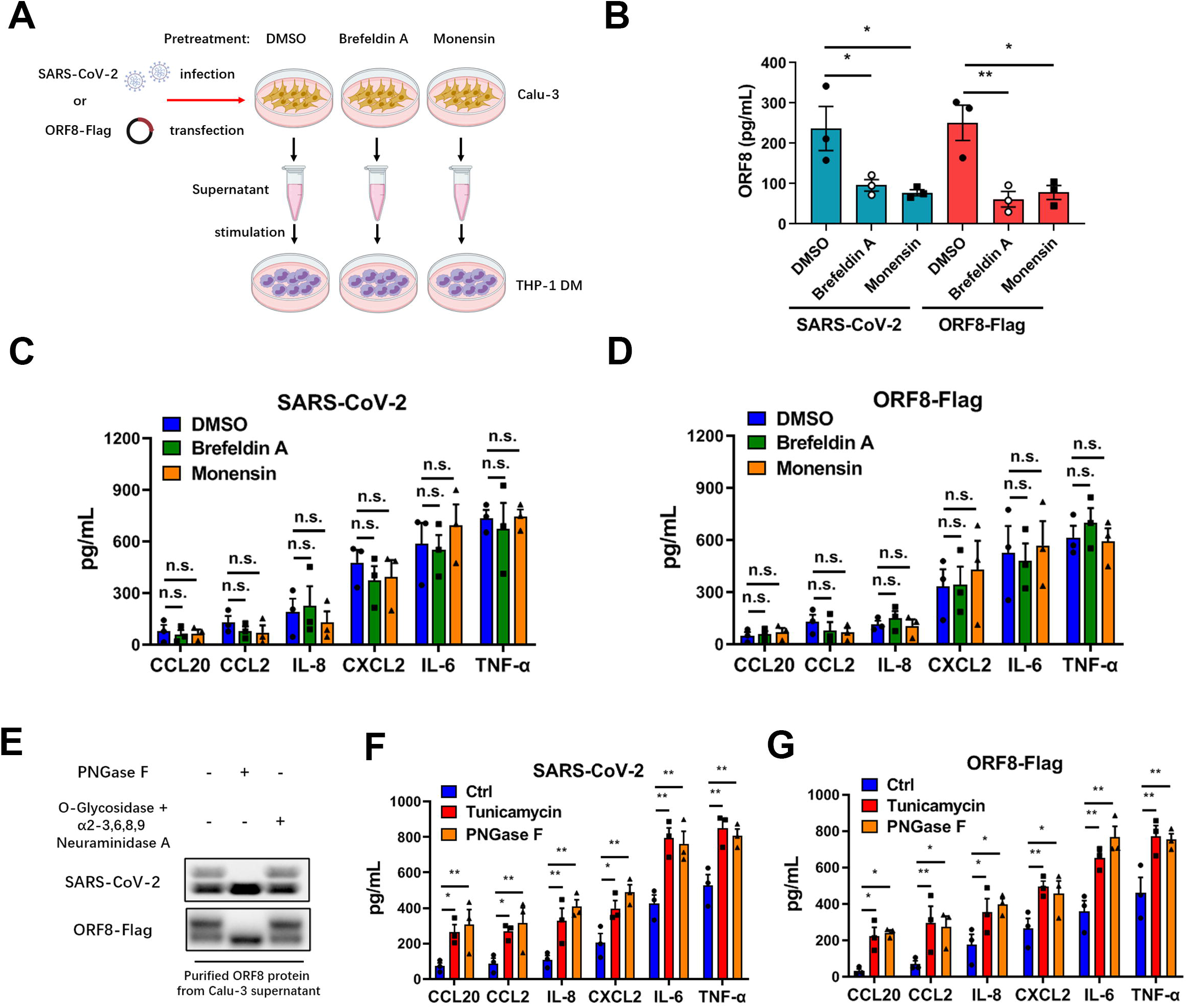
Unconventional secretion of ORF8 induces cytokine storm. (A) Schematic diagram of unconventional secretion model upon SARS-CoV-2 infection. Brefeldin A (3 μg/mL) or Monensin (2 μM) was used to pretreat Calu-3 cells for 2 hours, followed by SARS-CoV-2 infection for 12 hours at a dosage of 10^5^ TCID_50_/mL, or Flag-tagged ORF8 plasmid transfection for 12 hours. The supernatant was collected to stimulate THP-1 DM cells. (B) The secretion of ORF8 in (A) was detected by ELISA. Data are shown as the mean ± s.e.m. of n = 3 biological replicates. (C, D) The release of cytokines and chemokines in (A) was detected by ELISA. Data are shown as the mean ± s.e.m. of n = 3 biological replicates. (E) PNGase F (1,000 units/µg protein), O-Glycosidase (4,000 units/µg protein) or α2-3, 6, 8, 9 Neuraminidase A (4 units/μg glycoprotein) was added into the purified SARS-CoV-2 ORF8 protein to release glycans. Western blotting was used to detect the glycosylation of ORF8 protein. Representative images from n = 3 biological replicates are shown. (F, G) Calu-3 cells were infected with SARS-CoV-2 (F) or transfected with ORF-Flag plasmids (G). Tunicamycin (2µg/mL) was added into Calu-3 cells for 2 hours to prevent N-linked glycosylation; PNGase F (1,000 units/µg protein) was used to remove the N-linked glycosylation in purified ORF8 protein. After deglycosylation assays, the cell culture supernatant or purified ORF8 protein was used to stimulate THP-1 DM cells. After 12 hours, the release of cytokines and chemokines was detected by ELISA. Data are shown as the mean ± s.e.m. of n = 3 biological replicates. One-way ANOVA followed by Bonferroni post *hoc* test (B-D, F, G) was used for data analysis. *, p < 0.05, **, p < 0.01. Abbreviations: n.s., not significant.

The current data point to a potential possibility that ORF8 secreted via different pathways might undertake different responsibilities. Interestingly, in our western blots, a smear band shifted to a higher molecular mass compared to ORF8 was regularly observed, this smear however disappeared when the conventional secretory pathway was inhibited by signal peptide deletion or Golgi apparatus damage (Fig. 2B, D, E). Increasing the acrylamide concentration of the SDS-PAGE gel and prolonging the electrophoresis time, we were finally able to distinguish a second band of secreted SARS-CoV-2 ORF8 (Fig. 3E). The Golgi apparatus is known to be the workshop for protein trafficking and processing(*32*), the most common form of protein processing in Golgi apparatus is glycosylation(*16*). With this in mind, we asked whether ORF8 is glycosylated during the conventional secretory pathway, which might be responsible for the band shift. Using PNGase F or O-Glycosidase + α2-3, 6, 8, 9 Neuraminidase A, we tested the glycosylation status of ORF8 protein secreted from Calu-3 cells infected with SARS-CoV-2. We found that PNGase F treatment, which hydrolyzes most of the N-linked glycans(*33, 34*), was leading to the formation of a single ORF8 protein band (Fig. 3E). Digesting SARS-CoV-2 ORF8 with O-linked glycan hydrolase O-Glycosidase and α2-3, 6, 8, 9 Neuraminidase A did not change the type of bands compared to untreated samples (Fig. 3E). These data suggested that part of the secreted SARS-CoV-2 ORF8 protein was N-glycosylated, and that this is likely a result of conventional secretion through the Golgi apparatus. By contrast, SARS ORF8a did not respond to glycoside hydrolases at all (Fig. S2C), which means that conventional pathway secreted SARS ORF8a is not glycosylated.

Next, we asked whether the glycosylation status of ORF8 is associated with the release of pro-inflammatory factors. Calu-3 epithelial cells were infected with SARS-CoV-2, and tunicamycin(*35*) or PNGase F was used to oppose the N-linked glycosylation, respectively. Then purified ORF8 protein was added into the culture medium of THP-1 DM cells for stimulation. The results showed that macrophages stimulated with non-glycosylated ORF8 showed an elevated level of cytokine secretion (Fig. 3F). Furthermore, we used plasmid transfection instead of SARS-CoV-2 infection to validate this result. In line with previous results, a similar upregulation in cytokine release was observed in the tunicamycin and PNGase F treatment groups (Fig. 3G). In contrast, the pro-inflammatory factor expression of Calu-3 cells transfected with SARS ORF8a-Flag plasmid did not change upon tunicamycin or PNGase F treatment (Fig. S2D). Taken together, these results suggested that glycosylation state of ORF8 is closely associated with the release of pro-inflammatory factors.

### N-linked glycosylation at Asn78 impedes ORF8 binding to IL17RA

In order to determine the specific glycosylation site of SARS-CoV-2 ORF8, we collected the supernatant of SARS-CoV-2 infected Calu-3 epithelial cells and performed high performance liquid chromatography-tandem mass spectrometry (HPLC-MS/MS) for N-glycosylation site mapping (Fig. 4A). According to the identification by MS, we found the Asparagine 78 (Asn78 or N78) of ORF8 protein to be glycosylated (Fig. 4B). We further investigated the glycosylation site by creating a SARS-CoV-2 variant carrying the ORF8 N78Q mutation. Following infection with a SARS-CoV-2 ORF8-N78Q mutant, Calu-3 epithelial cells secreted a single form of unglycosylated ORF8 (Fig. 4C). This finding was also validated by transfection of exogenously expressed Flag-tagged N78Q ORF8 plasmid (Fig. 4C). Further, we treated THP-1 DM cells with supernatant enriched from Calu-3 cells infected with wild-type or the SARS-CoV-2 ORF8-N78Q mutant, respectively. In this context, more pronounced cytokine release was observed in macrophages stimulated with the ORF8-N78Q mutant protein (Fig. 4D). This result was further validated by exogenous expression of wild-type ORF8 and N78Q mutants. Consistent with previous results, transfection of N78Q mutant led to the excessive expression of pro-inflammatory factors (Fig. 4E). These data show that unglycosylated ORF8, instead of glycosylated ORF8, is involved in the inflammation response upon SARS-CoV-2 infection.

**Figure 4.**
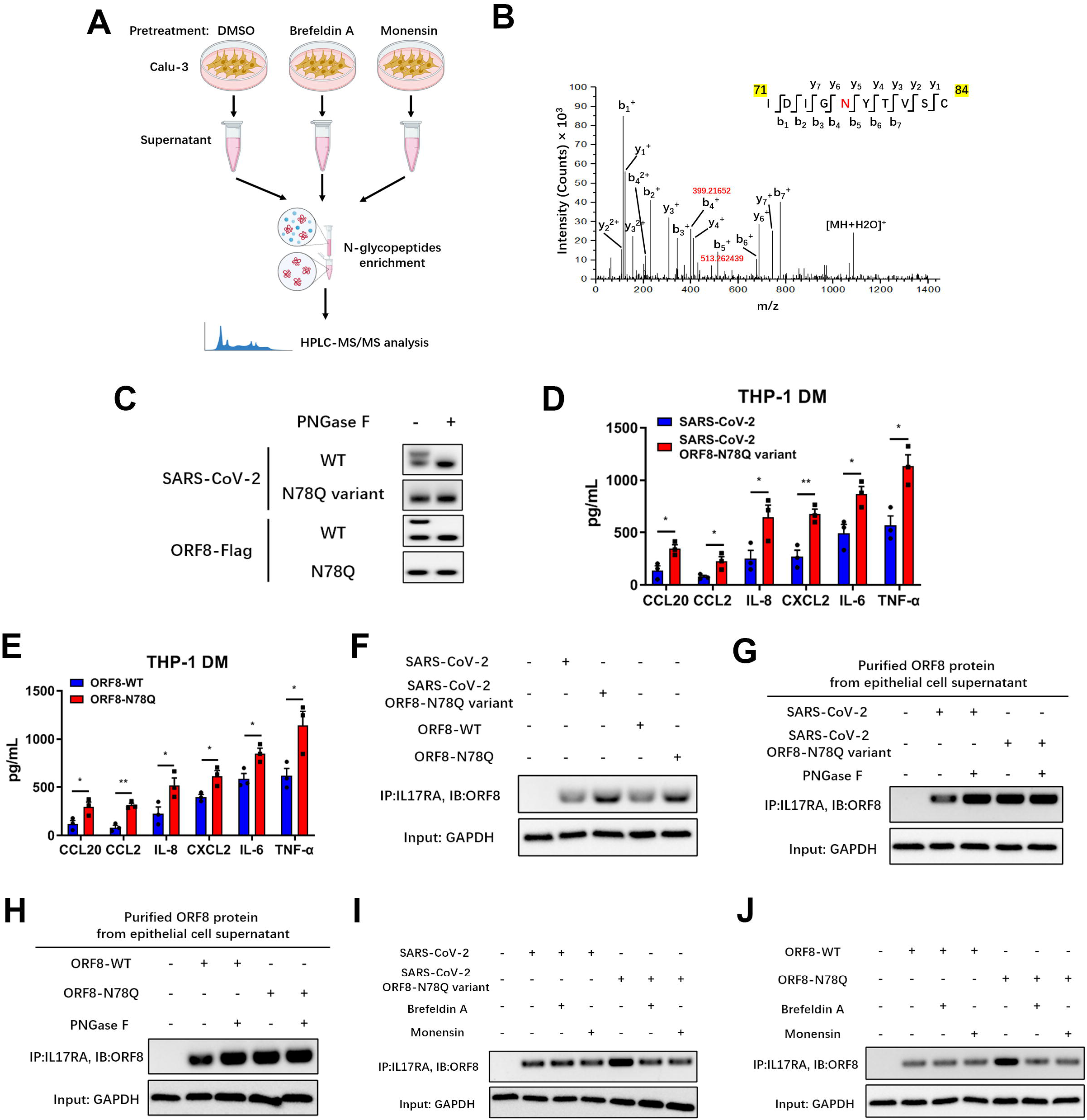
ORF8 N78 glycosylation blocks its interaction with IL17RA. (A) Schematic diagram of glycosylation identification based on HLPC-MS/MS. DMSO (control), Brefeldin A (3 μg/mL) or Monensin (2 μM) was used to pretreat Calu-3 cells for 2 hours. The supernatant was collected for HLPC-MS/MS analysis. (B) SARS-CoV-2 ORF8 secreted through conventional pattern has N78 glycosylation. An increase of 0.002989 Da of Asn residue was used to determine N-linked glycosylation. (C-E) Calu-3 cells were infected with SARS-CoV-2 variant carrying ORF8 N78Q mutant (C, D), or transfected with ORF8 N78Q plasmids (C, E). After 12 hours, the supernatant was collected and divided into two parts. One part was used to purify ORF8 protein, followed by PNGase F digestion and western blotting (C); the other part was used to stimulate THP-1 DM cells for 12 hours, followed by detection of cytokines and chemokines by ELISA (D, E). Representative images from n = 3 biological replicates are shown (C). Data are shown as the mean ± s.e.m. of n = 3 biological replicates (D, E). (F) Calu-3 cells were infected with SARS-CoV-2 ORF8-N78Q variant, or transfected with ORF8-N78Q plasmids. Twelve hours later, the supernatant was used to stimulate THP-1 DM cells for another 12 hours. The interaction of ORF8 and IL17RA was detected by co-immunoprecipitation. Representative images from n = 3 biological replicates are shown. (G, H) Calu-3 cells were infected with SARS-CoV-2 ORF8-N78Q variant (G), or transfected with ORF8-N78Q plasmids (H). After 12 hours, the supernatant was collected to purify ORF8 protein. After PNGase F digestion, the ORF8 protein was used to stimulate THP-1 DM cells. After 12 hours, the interaction of ORF8 and IL17RA was detected by co-immunoprecipitation. Representative images from n = 3 biological replicates are shown. (I, J) Brefeldin A (3 μg/mL) or Monensin (2 μM) was used to pretreat Calu-3 cells for 2 hours, followed by infection with SARS-CoV-2 ORF8-N78Q variant (I), or transfection with ORF8-N78Q plasmids (J). Twelve hours later, the supernatant was used to stimulate THP-1 DM cells for another 12 hours. The interaction of ORF8 and IL17RA was detected by co-immunoprecipitation. Representative images from n = 3 biological replicates are shown. One-way ANOVA followed by Bonferroni post *hoc* test (D, E) was used for data analysis. *, p < 0.05, **, p < 0.01.

In our previous studies, we found that the interaction between ORF8 and host IL17RA contributes to the formation of a cytokine storm(*6*). Here, we tested the effect of ORF8 glycosylation on the activation of the IL-17 pathway. We found that ORF8 protein secreted from the SARS-CoV-2 ORF8-N78Q variant or Flag-tagged ORF8-N78Q mutant, which could not be N-glycosylated at N78, exhibited stronger binding to the IL17RA receptor compared to a wild-type control (Fig. 4F). Further, the interaction between SARS-CoV-2 ORF8 and IL17RA was significantly increased when PNGase F was used to remove glycosylation (Fig. 4G). However, ORF8-N78Q showed consistently strong binding to IL17RA regardless of PNGase F treatment (Fig. 4G). We also tested the activation of NF-κB signaling downstream of the IL-17 pathway(*36*). Consistently, the activation of NF-κB signaling was positively correlated with the binding affinity between ORF8 and IL17RA (Fig. S3A, B). Data collected from plasmid transfection of Flag-tagged ORF8 mutant further validated this result (Fig. 4H, S3C). To further confirm that the N-linked glycosylated ORF8 was secreted via a conventional pathway, Brefeldin A or Monensin was used to pretreat Calu-3 epithelial cells to block conventional secretory transport. As a result, inhibition of Golgi-dependent vesicle transport decreased the interaction between ORF8 and IL17RA in N78Q groups, because less ORF8 was secreted when ER-Golgi trafficking was blocked (Fig. S3D, E); however, this inhibition did not affect the interaction between ORF8 and IL17RA in control groups, mainly due to conventionally secreted ORF8 was glycosylated (Fig. 4I, J, S3F, G).

Further, we prepared N-glycosylated ORF8 protein (ORF8-N-Glyc) *in vitro* and stimulated THP-1 DM cells directly. The results showed that glycosylated ORF8 could not bind the IL17RA receptor (Fig. 5A), and that secretion of pro-inflammatory factors was significantly reduced (Fig. 5B). These data indicated that glycosylation-deficient ORF8 is capable of binding to IL17RA and subsequent activation of the IL-17 pathway, thus promoting the cytokine storm. In order to further verify the contribution of ORF8 glycosylation to cytokine release during SARS-CoV-2 infection *in vivo*, we treated humanized ACE2 (hACE2) mice with aerosols of synthetic ORF8 or ORF8-N-Glyc. Compared with control group, the survival time of mice exposed to ORF8-N-Glyc was significantly prolonged (Fig. 5C). We also observed very little inflammation in the lungs from mice treated with ORF8-N-Glyc, while the lung lesions in unglycosylated ORF8-exposed mice were much more severe (Fig. 5D). Additionally, mice treated with ORF8-N-Glyc secreted decreased levels of cytokines and chemokines in lungs and livers (Fig. 5E, F). Taken together, these data indicated that the N78 glycosylation of ORF8 participates in the regulation of inflammatory response during SARS-CoV-2 infection.

**Figure 5.**
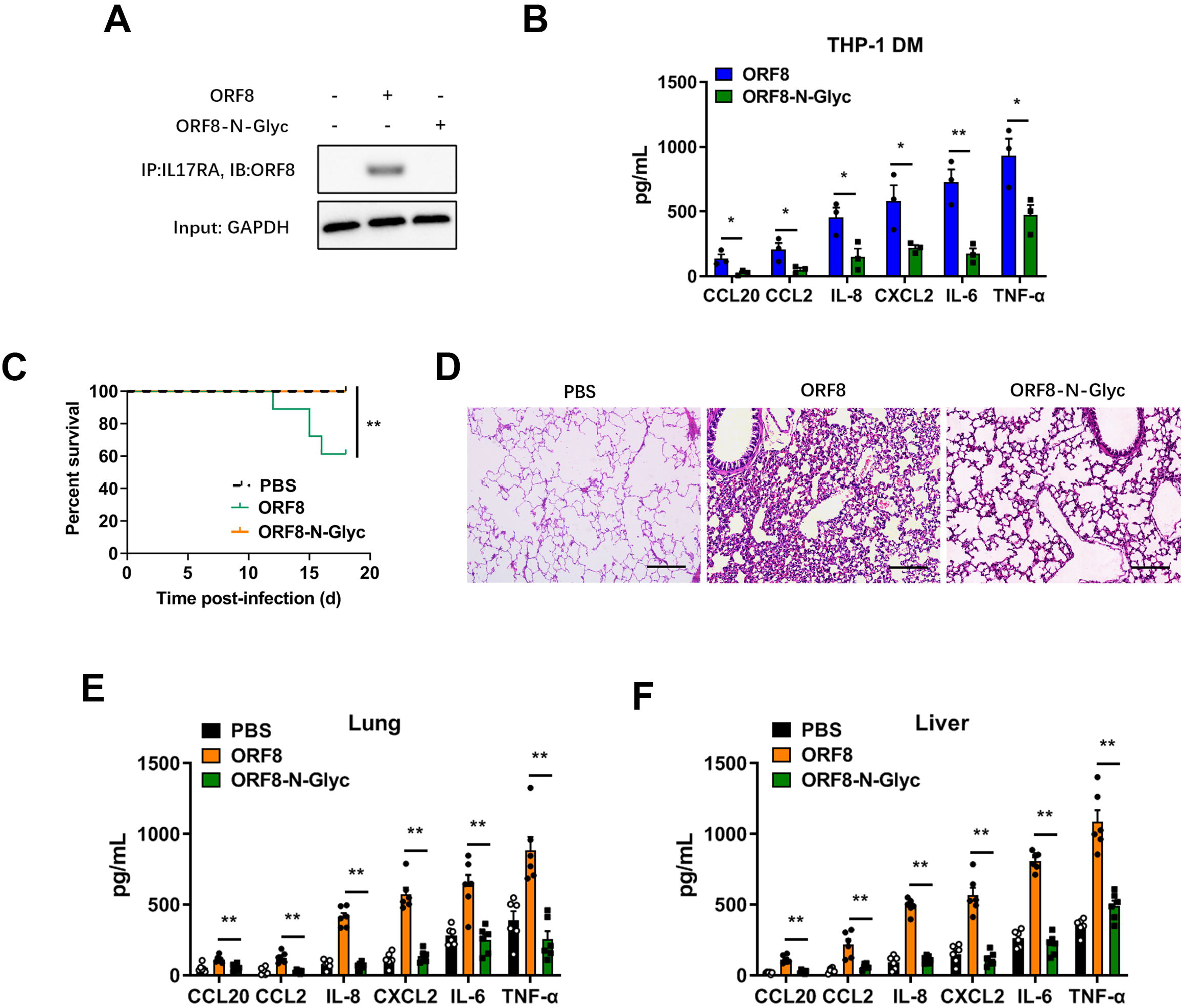
N-linked glycosylation of ORF8 protects mice from cytokine storm. (A, B) Synthetic N-linked-glycosylated ORF8 (ORF8-N-Glyc) (1 μg/mL) was added into the cell culture medium of THP-1 DM cells to stimulate IL-17 pathway. The interaction of ORF8 and IL17RA was detected by co-immunoprecipitation (A). The release of cytokines and chemokines was detected by ELISA (B). Representative images from n = 3 biological replicates are shown (A). Data are shown as the mean ± s.e.m. of n = 3 biological replicates (B). (C) Survival of K18-hACE2 mice infected with PBS (n = 11), unglycosylated ORF8 (n = 18) or synthetic N-linked-glycosylated ORF8 proteins (n = 17) (200 μg/mouse). Data are shown as Kaplan-Meier curves. (D) Lung lesions of K18-hACE2 mice infected with unglycosylated ORF8 or synthetic N-linked-glycosylated ORF8 proteins (200 μg/mouse) were detected by H&E staining at day 7 post infection (dpi). Representative images from n = 6 biological replicates are shown. Scale bar = 500 μm. (E, F) The release of cytokines and chemokines in lungs (E) and livers (F), which were obtained from K18-hACE2 mice intranasally infected with unglycosylated ORF8 or synthetic N-linked-glycosylated ORF8 proteins (200 μg/mouse), was detected by ELISA at 7 dpi. Data are shown as the mean ± s.e.m. of n = 6 biological replicates. Log-rank (Mantel-Cox) test (C) and one-way ANOVA followed by Bonferroni post *hoc* test (B, E, F) were used for data analysis. *, p < 0.05, **, p < 0.01.

### YIF1B is essential for the unconventional secretion of ORF8

A substantial number of secreted eukaryotic proteins lacking classical signal peptides (called leaderless cargoes) are released through unconventional secretion(*37, 38*). It has been reported that channel proteins located on the ER-Golgi intermediate compartment (ERGIC) might mediate translocation of leaderless cargoes into transport vesicles(*39*). However, the driving factors of initial vesicle formation have not yet been fully elucidated. Recent evidence suggests that autophagy contributes to the formation of unconventional secretory vesicles(*39–41*). Therefore, we next examined whether the unconventional secretion of ORF8 is regulated by autophagy. Calu-3 epithelial cells were pretreated with Brefeldin A to block the conventional secretory pathway, and then infected with SARS-CoV-2. Under the stimulation of starvation or Rapamycin (an autophagy activator), the secretion of ORF8 increased significantly (Fig. S4A). Autophagy inhibitor 3-Methyladenine (3-MA) or Wortmannin (Wtm) could counteract the starvation-induced ORF8 secretion (Fig. S4B). Consistently, knockdown of autophagy-related genes, such as *Atg5*, *Atg2a* or *Atg2b*, also inhibited the release of ORF8 (Fig. S4C). Additionally, we observed co-localization of ORF8 and ERGIC-53, indicating the possibility of ORF8 translocation into ERGIC (Fig. S4D). The proteinase K protection test also provided evidence of vesicle-mediated ORF8 transportation (Fig. S4E). These data suggested that SARS-CoV-2 ORF8 likely features an unconventional secretion pattern similar to eukaryotic leaderless cargoes.

To further determine the channel protein that guides the translocation of ORF8 into an unconventional secretion pattern, we performed mass spectrometry analysis to identify proteins that interact with unglycosylated ORF8. To enrich as many potential channel proteins on ORF8 transport vesicles as possible, dihydrofolate reductase (DHFR)-tagged ORF8 was transfected into HEK-293FT cells pretreated with aminopterin and Brefeldin A/Monensin as previously described (*12*)(Fig. S4F). Of the 156 potential interacting proteins, seven channel proteins appeared to be related to cargo transport (Fig. S4G). We then screened the seven candidates using knockdown strategy by siRNAs, and found that YIF1B was associated with the unconventional secretion of ORF8 (Fig. S4G).

Next, we verified the interaction between ORF8 and YIF1B by immunoprecipitation. Flag-tagged ORF8 was transfected into HEK-293FT cells and an interaction between ORF8 and YIF1B was observed (Fig. 6A). The co-localization of exogenously expressed HA-tagged YIF1B and Flag-tagged ORF8 was detected in HEK-293FT cells by immunoprecipitation (Fig. 6B) and the Duolink proximity ligation assay (Fig. 6C). Further, endogenous YIF1B and virus-derived ORF8 were able to form a complex in SARS-CoV-2 infected Calu-3 epithelial cells (Fig. 6D).

**Figure 6.**
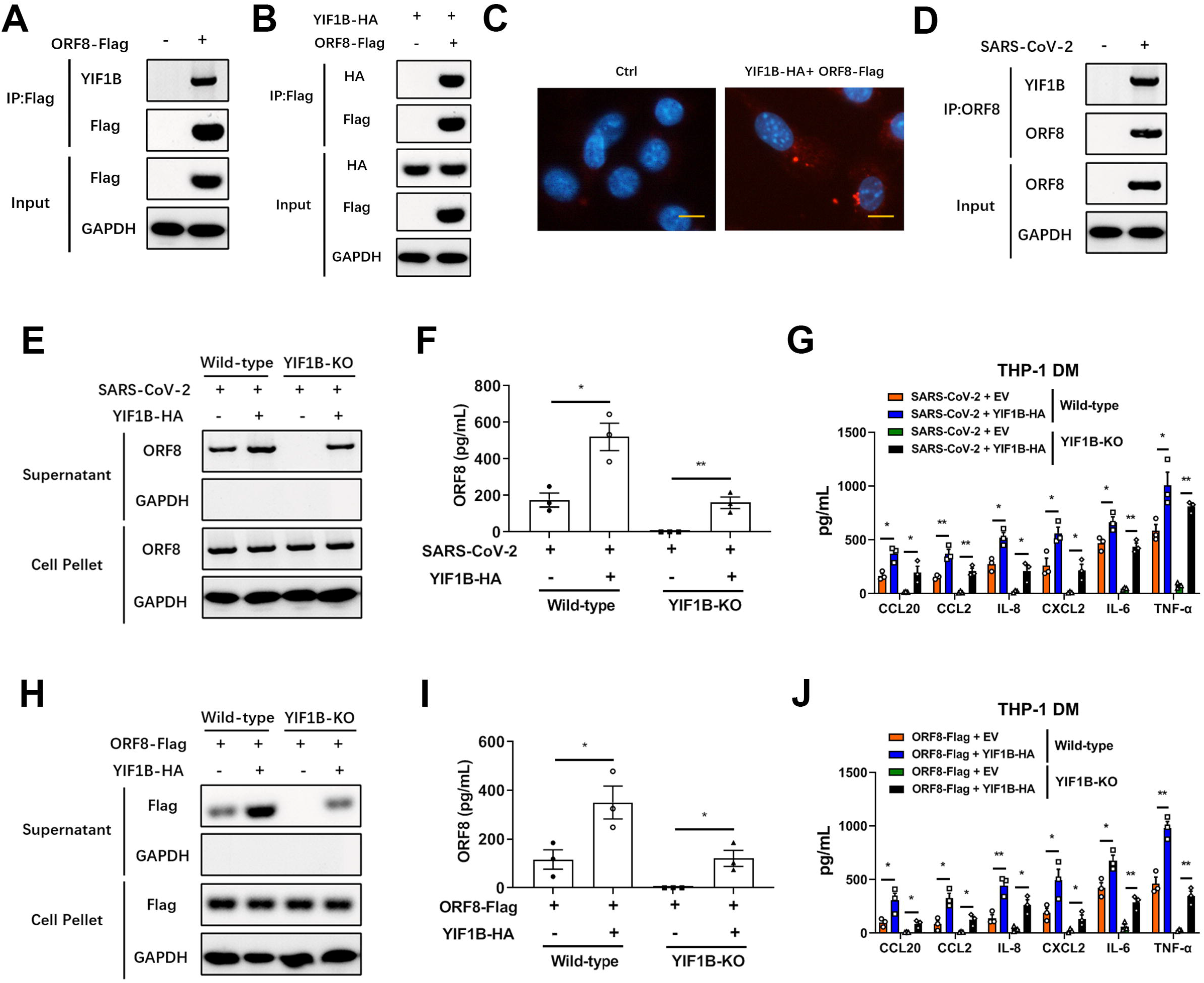
YIF1B interacts with unconventionally secreted ORF8. (A) Flag-tagged ORF8 was transfected into HEK-293FT cells. After 12 hours, the interaction of ORF8 and endogenic YIF1B was detected by co-immunoprecipitation. Representative images from n = 3 biological replicates are shown. (B, C) Flag-tagged ORF8 and HA-tagged YIF1B were co-transfected into HEK-293FT cells. After 12 hours, the interaction of ORF8 and YIF1B was detected by co-immunoprecipitation (B) and Duolink PLA assay (C). Representative images from n = 3 biological replicates are shown. (D) Calu-3 cells were infected with SARS-CoV-2 for 12 hours at a dosage of 10^5^ TCID_50_/mL. The interaction of ORF8 and YIF1B was detected by co-immunoprecipitation. Representative images from n = 3 biological replicates are shown. Scale bar = 10 μm. (E, F) Wild-type and YIF1B-KO cells were transfected with HA-tagged YIF1B, followed by infection with SARS-CoV-2 for 12 hours at a dosage of 10^5^ TCID_50_/mL. The secretion of ORF8 was detected by western blotting (E) and ELISA (F). Brefeldin A (3 μg/mL) was used to block conventional secretion of ORF8. Representative images from n = 3 biological replicates are shown (E). Data are shown as the mean ± s.e.m. of n = 3 biological replicates (F). (G) Wild-type and YIF1B-KO cells were transfected with HA-tagged YIF1B, and were infected with SARS-CoV-2 for 12 hours at a dosage of 10^5^ TCID_50_/mL. The cell culture supernatant was collected to stimulate THP-1 DM cells for 12 hours. The release of cytokines and chemokines was detected by ELISA. Data are shown as the mean ± s.e.m. of n = 3 biological replicates. (H, I) Wild-type and YIF1B-KO cells were co-transfected with Flag-tagged ORF8 and HA-tagged YIF1B. After12 hours, the secretion of ORF8 was detected by western blotting (H) and ELISA (I). Brefeldin A (3 μg/mL) was used to block conventional secretion of ORF8. Representative images from n = 3 biological replicates are shown (H). Data are shown as the mean ± s.e.m. of n = 3 biological replicates (I). (J) Wild-type and YIF1B-KO cells were co-transfected with Flag-tagged ORF8 and HA-tagged YIF1B. Twelve hours later, the cell culture supernatant was collected to stimulate THP-1 DM cells for another 12 hours. The release of cytokines and chemokines was detected by ELISA. Data are shown as the mean ± s.e.m. of n = 3 biological replicates. Unpaired two-tailed Student *t* test (F, I) and one-way ANOVA followed by Bonferroni post *hoc* test (G, J) were used for data analysis. *, p < 0.05, **, p < 0.01.

Then we generated YIF1B-deficient cells (YIF1B-KO) based on the Calu-3 cell line, and found that unconventional secretion of ORF8 disappeared in absence of YIF1B; however, exogenous supplementation of YIF1B by plasmid transfection rescued the secretion of ORF8 (Fig. 6E, F). Moreover, cytokine release assays further proved that YIF1B overexpression reconstituted the function of ORF8 protein in YIF1B-KO cells (Fig. 6G). Likewise, ORF8 secretion and cytokine release data collected from ORF8-Flag plasmid transfection further confirmed above findings (Fig. 6H-J). These results indicated that the unconventional secretion pattern of SARS-CoV-2 ORF8 requires host YIF1B.

### The α4 helix of YIF1B recognizes the β8 sheet of ORF8 for interaction

The structural domain of ORF8 protein, which is eight β-pleated sheets consisted by 165 amino acid chain, has been reported(*42*) (Fig. 7A). We constructed ORF8 Δβ1-Δβ8 deletion mutants, respectively. The results of co-immunoprecipitation showed that an ORF8 mutant lacking the β8 sheet could not bind to YIF1B (Fig. 7B). Similarly, we observed that β8 sheet-deleted ORF8 could not be secreted when the conventional secretion pathway was blocked by Brefeldin A pretreatment (Fig. 7C, D). Likewise, the cytokine release by THP-1 DM cells stimulated with supernatant proved that β8 sheet domain was closely associated with the unconventional secretory pathway of ORF8 protein (Fig. 7E).

**Figure 7.**
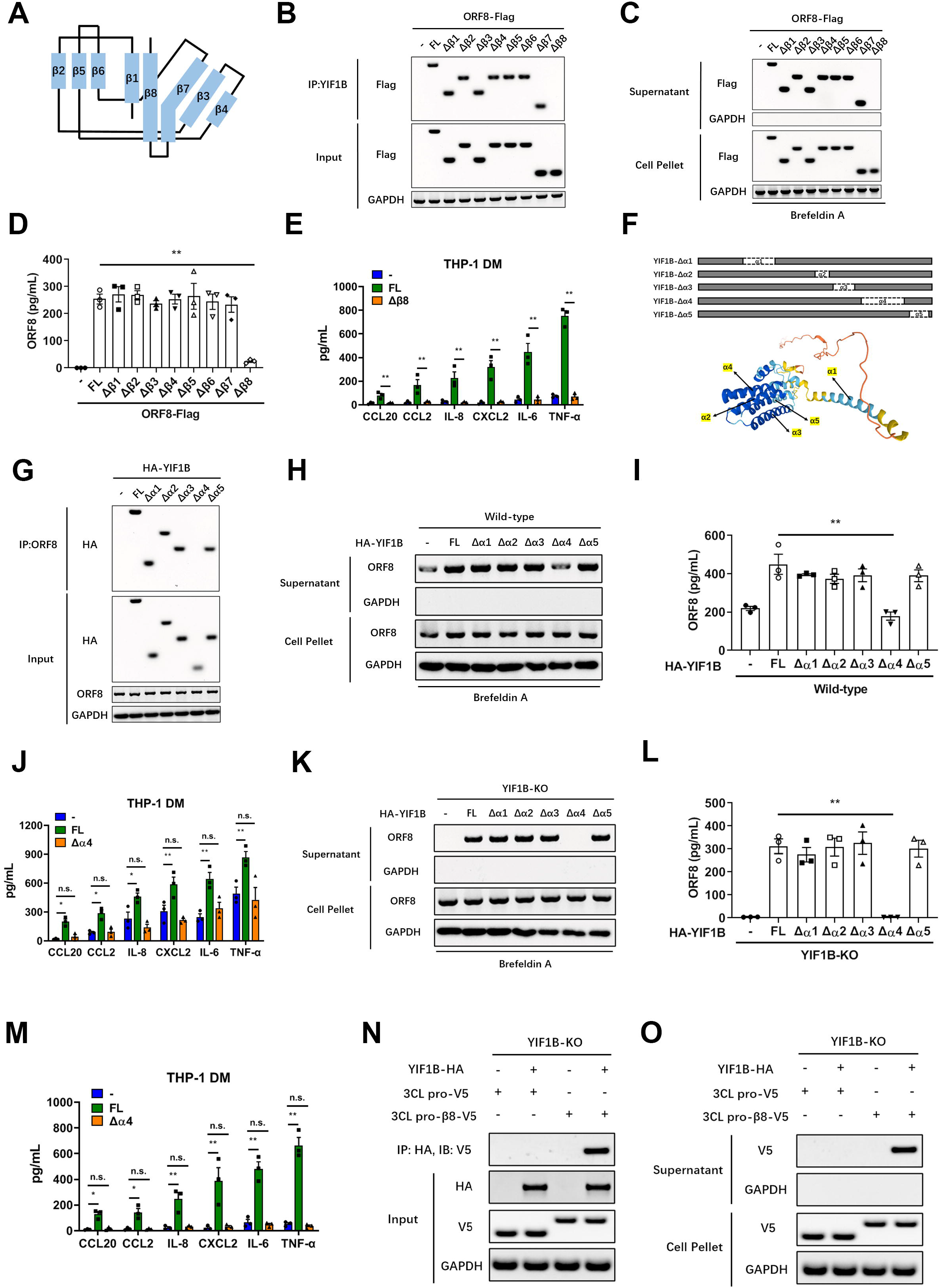
β8 sheet of ORF8 and α4 helix of YIF1B are essential for the interaction. (A) Structure of β-pleated sheet in SARS-CoV-2 ORF8 protein. (B) ORF8 β sheet mutants (Δβ1-Δβ8) were transfected into Calu-3 cells. After 12 hours, the interaction of ORF8 and YIF1B was detected by co-immunoprecipitation. Representative images from n = 3 biological replicates are shown. (C, D) ORF8 β sheet mutants (Δβ1-Δβ8) were transfected into Calu-3 cells. After 12 hours, the secretion of ORF8 was detected by western blotting (C) and ELISA (D). Brefeldin A (3 μg/mL) was used to block conventional secretion of ORF8. Representative images from n = 3 biological replicates are shown (C). Data are shown as the mean ± s.e.m. of n = 3 biological replicates (D). (E) ORF8 Δβ8 mutant was transfected into Calu-3 cells. Twelve hours later, the cell culture supernatant was collected to stimulate THP-1 DM cells for another 12 hours. The release of cytokines and chemokines was detected by ELISA. Brefeldin A (3 μg/mL) was used to block conventional secretion of ORF8. Data are shown as the mean ± s.e.m. of n = 3 biological replicates. (F) Structure of α helix and corresponding deletion mutants of YIF1B protein. (G) YIF1B α helix mutants (Δα1-Δα5) were transfected into Calu-3 cells. After 12 hours, the interaction of ORF8 and YIF1B was detected by co-immunoprecipitation. Representative images from n = 3 biological replicates are shown. (H, I) YIF1B α-helix mutants (Δα1-Δα5) were transfected into wild-type Calu-3 cells. After 12 hours, the secretion of ORF8 was detected by western blotting (H) and ELISA (I). Brefeldin A (3 μg/mL) was used to block conventional secretion of ORF8. Representative images from n = 3 biological replicates are shown (H). Data are shown as the mean ± s.e.m. of n = 3 biological replicates (I). (J) YIF1B Δα4 mutant was transfected into wild-type Calu-3 cells. Twelve hours later, the cell culture supernatant was collected to stimulate THP-1 DM cells for another 12 hours. The release of cytokines and chemokines was detected by ELISA. Brefeldin A (3 μg/mL) was used to block conventional secretion of ORF8. Data are shown as the mean ± s.e.m. of n = 3 biological replicates. (K, L) YIF1B α-helix mutants (Δα1-Δα5) were transfected into YIF1B-KO Calu-3 cells. After 12 hours, the secretion of ORF8 was detected by western blotting (K) and ELISA (L). Brefeldin A (3 μg/mL) was used to block conventional secretion of ORF8. Representative images from n = 3 biological replicates are shown (K). Data are shown as the mean ± s.e.m. of n = 3 biological replicates (L). (M) YIF1B Δα4 mutant was transfected into YIF1B-KO Calu-3 cells. The cell culture supernatant was collected to stimulate THP-1 DM cells for 12 hours. The release of cytokines and chemokines was detected by ELISA. Brefeldin A (3 μg/mL) was used to block conventional secretion of ORF8. Data are shown as the mean ± s.e.m. of n = 3 biological replicates. (N, O) HA-tagged YIF1B, V5-tagged 3CL pro and V5-tagged 3CL pro containing β8 sheet were transfected into YIF1B-KO Calu-3 cells. After 12 hours later, the interaction of YIF1B and 3CL pro was detected by co-immunoprecipitation (N). The secretion of 3CL pro was detected by western blotting (O). Representative images from n = 3 biological replicates are shown. One-way ANOVA followed by Bonferroni post *hoc* test (E, I, J, L, M) was used for data analysis. *, p < 0.05, **, p < 0.01. Abbreviations: n.s., not significant.

In order to understand how YIF1B mediates the recognition and secretion of leadless cargoes, we constructed deletion mutants of five α helixes located on the transmembrane domains of YIF1B according to the structural predication (https://alphafold.ebi.ac.uk/entry/Q5BJH7) (Fig. 7F). Co-immunoprecipitation results showed that a α4 helix deletion impaired the interaction between YIF1B and ORF8 (Fig. 7G). Next, we transfected exogenous YIF1B mutants into wild-type epithelial Calu-3 cells. We found that the unconventional secretion of ORF8 increased significantly in YIF1B mutants groups; this however was not observed in the Δα4 mutant group (Fig. 7H-J). Further, after transfection of YIF1B mutants into YIF1B-KO cells, the unconventional secretion of ORF8 was rescued by all YIF1B mutants, except the Δα4 mutant (Fig. 7K-M). These data implied that the unconventional secretion of ORF8 relies on the α4 helix of YIF1B and depends on its presence in a dose-dependent manner (Fig. 7H-J).

Next, we asked whether YIF1B-guided unconventional secretion of leaderless ORF8 is driven by direct recognition of the β8 sheet. To answer this question, we constructed a 3CL pro mutant with a β8 sheet fused to the C-terminus. Interestingly, an interaction between 3CL pro-β8 fusion and YIF1B was observed (Fig. 7N), and secreted 3CL pro could be collected from culture supernatant (Fig. 7O). Collectively, these evidences indicated that unconventional secretion of ORF8 is mediated by host YIF1B through recognition and interaction between the β8 sheet and α4 helix domain.

### YIF1B directly promotes the translocation of ORF8

Since the interaction between ORF8 and YIF1B is important for the unconventional secretion pattern of ORF8, we first looked for intracellular co-localization of ORF8 and YIF1B during SARS-CoV-2 infection. Immunofluorescence results showed that full-length ORF8 and YIF1B did co-localized (Fig. 8A). However, neither an ORF8 β8 sheet-deleted mutant nor a YIF1B α4 helix-deficient mutant co-localized with their respective partner (Fig. 8B, C). Next, a proteinase K protection test was performed to examine whether ORF8 transportation into vesicles relies on ORF8-YIF1B interaction. Full-length ORF8-Flag or YIF1B truncations were transfected into YIF1B-KO cells as indicated, the membrane pellets were collected by centrifugation and treated with proteinase K or Triton X-100. The results showed that in YIF1B-deficient cells, ORF8 was not transported into vesicles, leading to its degradation by proteinase K (Fig. 8D). On the contrary, ORF8 was able to resist the degradation by proteinase K when YIF1B truncation was counteracted by transfection of YIF1B plasmids, except in case of the Δα4 mutant (Fig. 8D).

**Figure 8.**
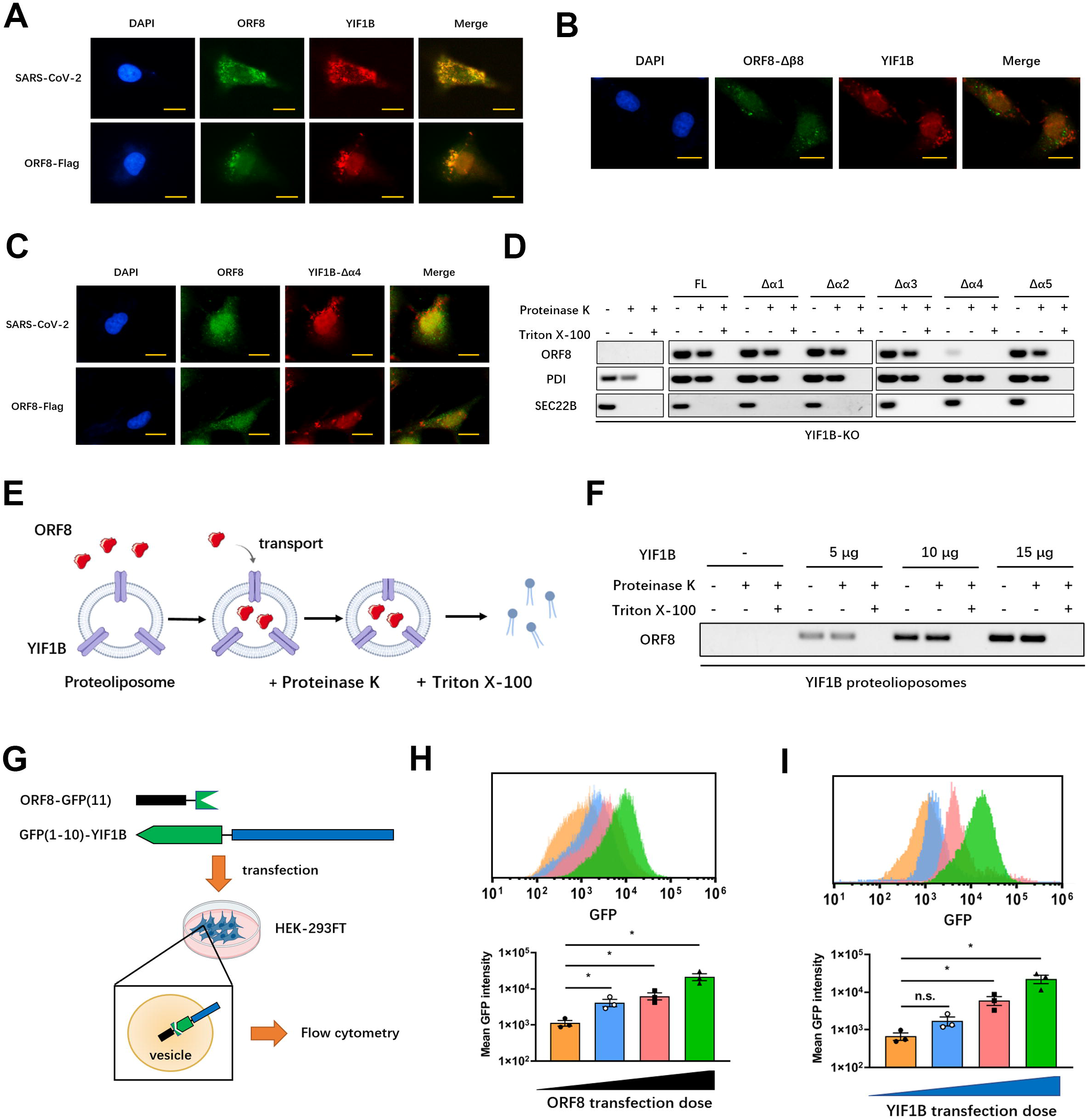
YIF1B promotes the transport of ORF8. (A) Calu-3 cells were infected with SARS-CoV-2, or transfected with ORF8-Flag. After 12 hours, the co-localization of ORF8 and YIF1B was observed by immunofluorescence. Representative images from n = 3 biological replicates are shown. Scale bar = 10 μm. (B) Calu-3 cells were transfected with ORF8-Δβ8 mutant. After 12 hours, the co-localization of ORF8 and YIF1B was observed by immunofluorescence. Representative images from n = 3 biological replicates are shown. Scale bar = 10 μm. (C) Calu-3 cells were infected with SARS-CoV-2, or transfected with ORF8-Flag. YIF1B-Δα4 mutant was also transfected into Calu-3 cells. After 12 hours, the co-localization of ORF8 and YIF1B was observed by immunofluorescence. Representative images from n = 3 biological replicates are shown. Scale bar = 10 μm. (D) ORF8 and YIF1B α-helix mutants (Δα1-Δα5) were transfected into YIF1B-KO Calu-3 cells. Whole cell lysates were collected for Proteinase K protection assay. PDI and SEC22B were used as positive control and negative control, respectively. Representative images from n = 3 biological replicates are shown. (E, F) Schematic diagram of *in vitro* translocation assay (E). Different doses of YIF1B protein and lipids were assembled to proteoliposomes. ORF8 protein and synthetic proteoliposomes were mixed in the lysate of HEK-293FT cells (without endomembrane). Proteoliposomes containing ORF8 were collected and aliquoted into three parts for Proteinase K protection assay (F). Representative images from n = 3 biological replicates are shown (F). (G) Schematic diagram of GFP fluorescence complementation system. ORF8 fused with GFP (11) and YIF1B containing GFP (1-10), were co-transfected into HEK-293FT cells, and flow cytometry was used to measure GFP fluorescence intensity. (H, I) Different doses of ORF8-GFP(11) were co-transfected with YIF1B-GFP(1-10) (H), or different doses of YIF1B-GFP(1-10) (I) were co-transfected with ORF8-GFP(11) into HEK-293FT cells, and flow cytometry was used to measure GFP fluorescence intensity. Representative images from n = 3 biological replicates are shown. Data are shown as the mean ± s.e.m. of n = 3 biological replicates. One-way ANOVA followed by Bonferroni post *hoc* test (H, I) was used for data analysis. *, p < 0.05. Abbreviations: n.s., not significant.

Furthermore, we constructed an *in vitro* ORF8 transport system as previously described(*12*). In this system, synthesized ORF8 was mixed with assembled proteoliposomes, and HEK-293FT cell lysate without endomembrane was used as the reaction buffer to promote the transport of ORF8 (Fig. 8E). A proteinase K protection test showed that YIF1B mediated the transport of ORF8 into proteoliposomes in a dose-dependent manner (Fig. 8F).

We also established a GFP fluorescence complementation system(*43*) to validate if ORF8 resides in the same vesicle that also harbors YIF1B. ORF8 fused with GFP(11) and YIF1B containing GFP(1–10) were co-transfected into HEK-293FT cells and flow cytometry was used to measure the fluorescence emitted by the combination of ORF8 and YIF1B (Fig. 8G). When the amount of transfected ORF8 or YIF1B increased, an elevation in fluorescence intensity was observed, suggesting that the binding rate of ORF8-YIF1B complex increased in a concentration depended manner(Fig. 8H, I). Collectively, these data clarified that YIF1B directly promotes the translocation of ORF8 into vesicles for unconventional secretion.

### YIF1B regulates cytokine storm upon SARS-CoV-2 infection

YIF1B regulates the translocation of SARS-CoV-2 ORF8 into vesicles, and thereby promotes the unconventional secretion of unglycosylated ORF8 protein, which is required for the downstream IL-17 signaling activation and release of pro-inflammatory. To ascertain the biologic significance of our findings, we generated YIF1B-deficient (*Yif1b*^-/-^) mice in a hACE2 background (Fig. S5A-C). After SARS-CoV-2 infection, the interaction between ORF8 and YIF1B disappeared (Fig. 9A), and ORF8 secretion was significantly decreased in alveolar epithelial cells collected from these mice (Fig. 9B). In line with our expectations, supplementation of exogenous YIF1B restored the secretion of ORF8 (Fig. 9B). Further, epithelial cells obtained from *Yif1b*^-/-^ mice only secreted N-glycosylated ORF8, indicating a deficiency of the unconventional secretory pathway (Fig. 9C). Consistently, ORF8 secreted from *Yif1b*^-/-^ epithelial cells neither bound the IL17RA receptor nor did it trigger the downstream NF-κB pathway (Fig. 9D, E). These data again illustrated that the unconventional secretory pathway mediated by YIF1B allows ORF8 to escape host cell glycosylation, thus enabling activation of the IL-17 signaling pathway.

**Figure 9.**
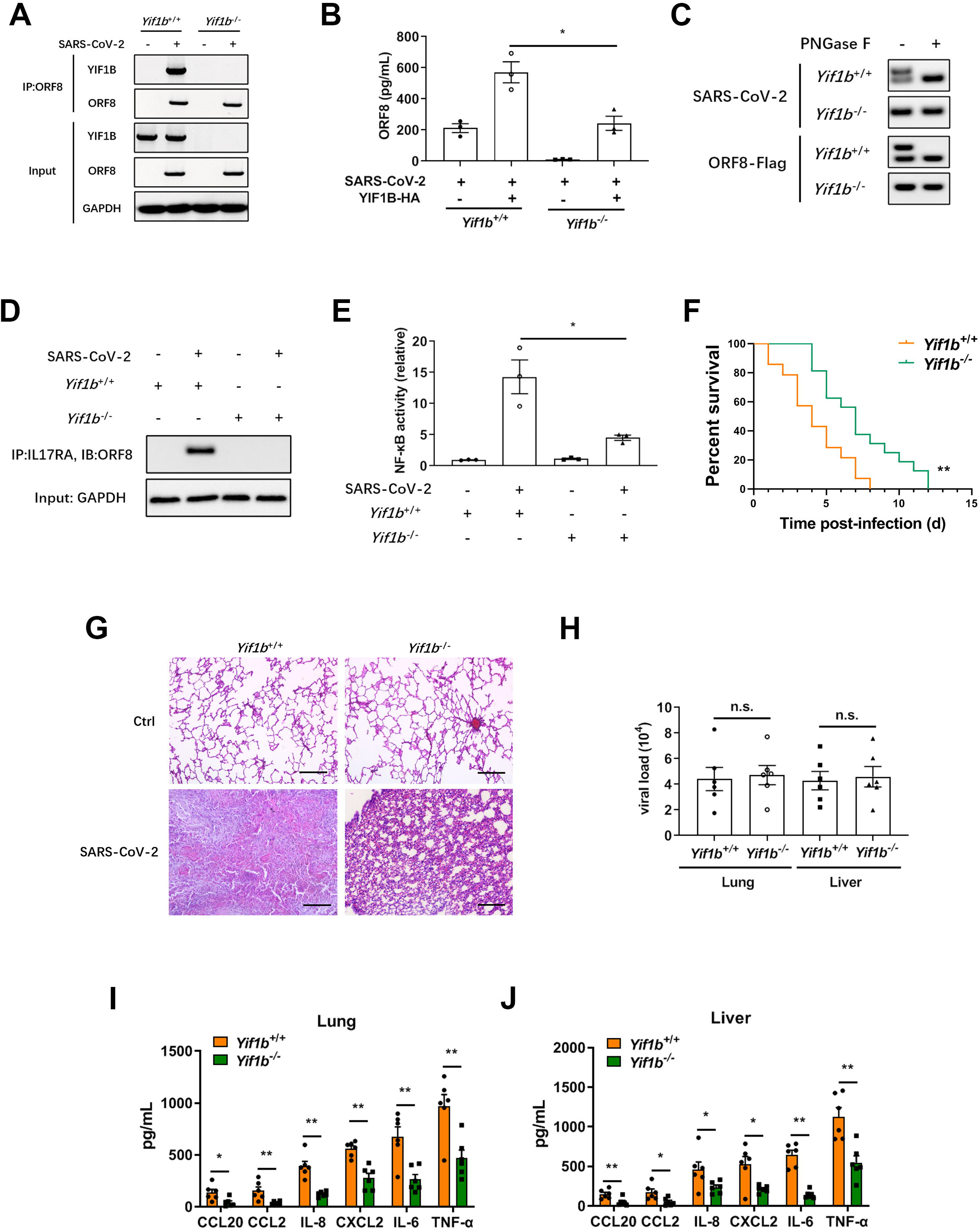
YIF1B-deficient mice resist to SARS-CoV-2-induced cytokine storm. (A) AECs obtained from *Yif1b*^-/-^ mice and their littermates (*Yif1b*^+/+^) were infected with SARS-CoV-2 for 12 hours at a dosage of 10^5^ TCID_50_/mL. The interaction of ORF8 and YIF1B was detected by co-immunoprecipitation (A). HA-tagged YIF1B was transfected into cells to restore YIF1B expression. The secretion of ORF8 was detected by ELISA (B). Representative images from n = 3 biological replicates are shown (A). Data are shown as the mean ± s.e.m. of n = 3 biological replicates (B). (C) AECs obtained from *Yif1b*^-/-^ mice and their littermates (*Yif1b*^+/+^) were infected with SARS-CoV-2, or transfected with ORF8-Flag plasmids. After 12 hours, the supernatant was collected to purify ORF8 protein. After PNGase F digestion, ORF8 protein was detected by western blotting. Representative images from n = 3 biological replicates are shown. (D, E) AECs obtained from *Yif1b*^-/-^ mice and their littermates (*Yif1b*^+/+^) were infected with SARS-CoV-2 at a dosage of 10^5^ TCID_50_/mL. After 12 hours, the supernatant was collected to purify ORF8 protein, which was used to stimulate THP-1 DM cells for another 12 hours. The interaction of ORF8 and IL17RA was detected by co-immunoprecipitation (D). The activation of IL-17 pathway was evaluated by testing NF-κB activity (E). Representative images from n = 3 biological replicates are shown (D). Data are shown as the mean ± s.e.m. of n = 3 biological replicates (E). (F) Survival of *Yif1b*^-/-^ mice (n = 16) and their littermates (*Yif1b*^+/+^, n = 14) after intranasal infection with SARS-CoV-2 with 4×10^5^ PFU. Data are shown as Kaplan-Meier curves. (G) Lung lesions of *Yif1b*^-/-^ mice and their littermates (*Yif1b*^+/+^) were detected by H&E staining at 7 dpi. Representative images from n = 6 biological replicates are shown. Scale bar = 500 μm. (H) Viral loads in lungs obtained from *Yif1b*^-/-^ mice and their littermates (*Yif1b*^+/+^) at 7 dpi. Data are shown as the mean ± s.e.m. of n = 6 biological replicates. (I, J) The secretion of cytokines and chemokines in lungs (I) and livers (J) obtained from *Yif1b*^-/-^ mice and their littermates (*Yif1b*^+/+^) at 7 dpi, was detected by ELISA. Data are shown as the mean ± s.e.m. of n = 6 biological replicates. Two-way ANOVA followed by Bonferroni post *hoc* test (B, E, I, J), Log-rank (Mantel-Cox) test (F) and unpaired two-tailed Student *t* test (H) were used for data analysis. *, p < 0.05, **, p < 0.01. Abbreviations: n.s., not significant.

To further characterize the functional role of host YIF1B in the development of a cytokine storm during SARS-CoV-2 infection, *Yif1b*^-/-^ hACE2 mice were intranasally infected with 4×10^5^ PFU plaque-forming units (PFU) of SARS-CoV-2. Compared to the *Yif1b*^+/+^ littermates, the survival time of *Yif1b*^-/-^ mice infected with SARS-CoV-2 was significantly prolonged (Fig. 9F). The lungs of *Yif1b*^-/-^ mice showed only mild inflammation compared to the extensive lung lesions observed in *Yif1b*^+/+^ mice (Fig. 9G). Further, we examined the viral loads and cytokine release in the spleens and livers of *Yif1b*^-/-^ mice and their *Yif1b*^+/+^ littermates. In this assay, although the viral loads showed no significant difference in the two groups (Fig. 9H), the cytokine release in spleens and livers of *Yif1b*^-/-^ mice was strongly alleviated compared to the control group (Fig. 9I, J). These results emphasize both the functional role of YIF1B in the unconventional secretion of ORF8 and the mechanistic role of unconventional ORF8 protein secretion in the development of a cytokine storm during SARS-CoV-2 infection.

Overall, we found that after invading host epithelial cells, SARS-CoV-2 ORF8 can be secreted through both conventional and conventional secretory pathways. In the conventional secretory pathway, ORF8 is N-glycosylated during the ER-Golgi trafficking, consequently, extracellular ORF8 lost the ability to recognize the IL17RA receptor of macrophages, likely due to steric hindrances imposed by N-glycosylation at the Asn78 site. By contrast, ORF8 is recognized and then translocated into vesicles directly by host YIF1B in an unconventional secretory pathway. Without experiencing the conventional ER-Golgi trafficking, ORF8 protein does not become glycosylated. Hence, extracellular ORF8 can be distributed through body fluid circulation and to get in contact with macrophages, where unglycosylated ORF8 binds the IL17RA receptor and activates the IL17 pathway and downstream NF-κ signaling facilitating the onset of a cytokine storm (Fig. 10).

**Figure 10.**
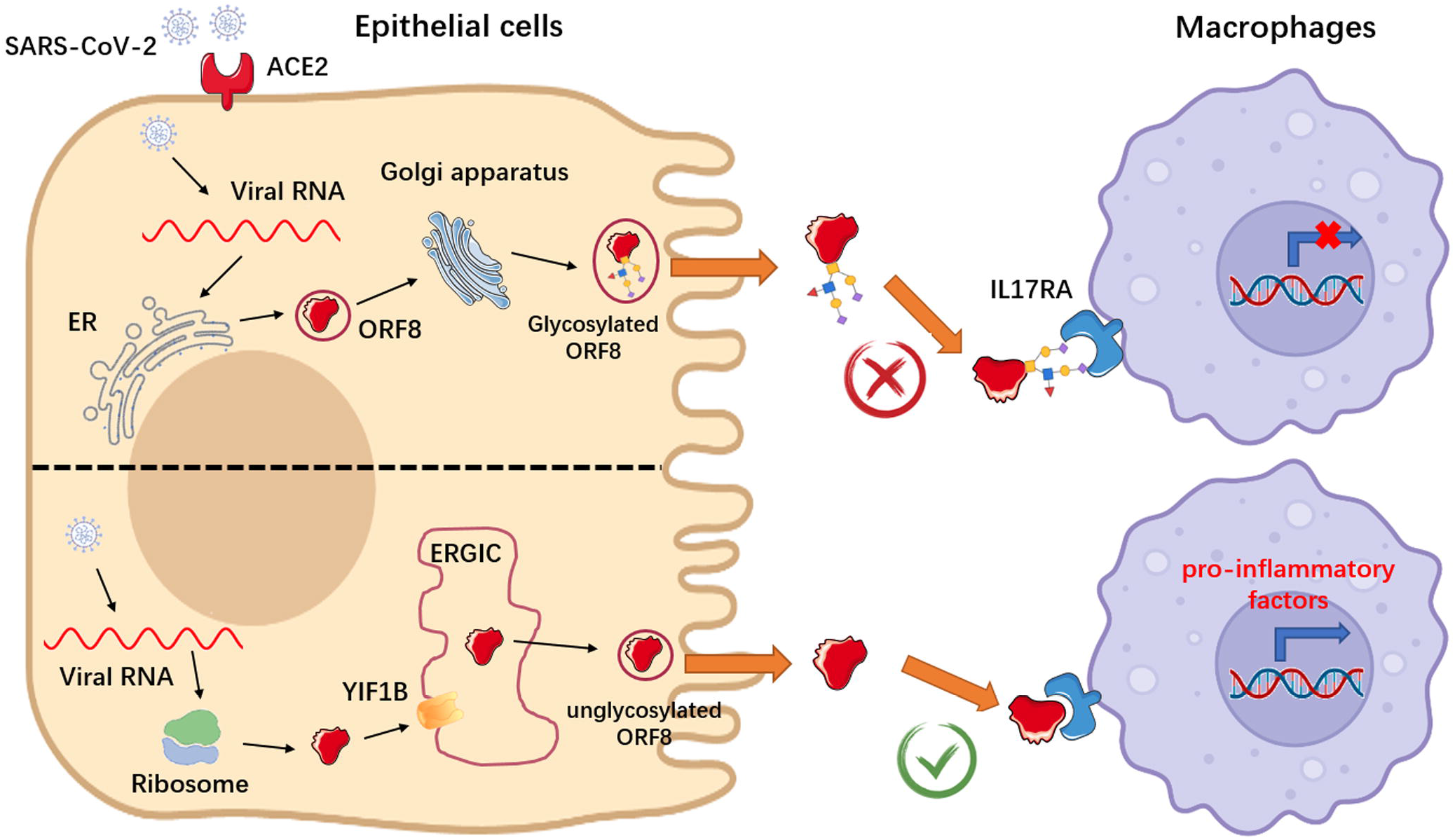
Schematic representation of conventional and unconventional secretion of ORF8 during SARS-CoV-2 infection.

## Discussion

Understanding the specific functions of SARS-CoV-2 proteins is pivotal for us to perceive the mechanisms which contribute to its high infectivity, fitness, and virulence. Numerous studies unravelling the differential functions of structural proteins have appeared recently(*28, 44*). However, it is worth noting that non-structural and accessory proteins encoded by SARS-CoV-2 likewise play significant roles in the regulation of the viral life cycle and also affect the immune response of the host. For example, ORF3a has been reported to induce apoptosis and promote lysosomal exocytosis-mediated viral egress(*45, 46*), while ORF6 protein of SARS-CoV-2 hampers the induction of host interferon signaling(*47*).

ORF8, a non-conserved accessory protein, is likely to be associated with the unique characteristics of SARS-CoV-2. According to clinical reports, ORF8 is highly immunogenic, anti-ORF8 antibodies are formed in the early stage of infection(*48*) and a significant T-cell response to ORF8 is observed in recovered patients(*49*). It is further reported that an ORF8-deficient SARS-CoV-2 strain (Δ382) in Singapore displays a significant reduction in virulence(*5*). In a recent study, Zhang et al. reported that SARS-CoV-2 ORF8 interacts with MHC-I, a marker protein located on the cell surface, and activates the lysosomal degradation pathway, thus achieving escape from immune surveillance by decreasing the expression of MHC I(*50*). These data indicate the specific role of ORF8 in infectivity and pathogenicity of SARS-CoV-2.

Our previous work showed that SARS-CoV-2 ORF8 could interact with the IL17RA receptor, thereby leading to IL-17 pathway activation and an increased secretion of pro-inflammatory factors(*6*). Considering that IL17RA is a transmembrane protein and ORF8 binds to the extracellular domain of IL17RA, as well as combining with the existing evidences, we proposed that ORF8 might be a secretory protein that is secreted into extracellular compartments. In this study, we show that SARS-CoV-2 ORF8 protein can be secreted by infected epithelial cells, which is supporting the role of ORF8 as a cellular messenger between alveolar epithelial cells and macrophages in the occurrence and development of a cytokine storm. Interestingly, we discovered that ORF8 can be secreted through both a conventional and an unconventional secretory pathway, and that ORF8 secreted via different pathways has different mechanistic functions in SARS-CoV-2 infection. Specifically, conventionally secreted ORF8 is N-glycosylated and loses the ability of binding the IL17RA receptor. Unconventionally secreted ORF8 is undergoing the conventional ER-Golgi trafficking pathway and consequently does not become glycosylated. Therefore, it is able to bind the IL17RA receptor and activate IL-17 signaling induce the expression of pro-inflammatory factors. The existence of this unique indirect cellular communication mechanism, instead of virion release, during the course of SARS-CoV-2 infection can at least partly explain why viral loads in patients are not directly proportional to the severity of disease symptoms in COVID-19(*51, 52*).

In this study, we found that SARS ORF8a is also a secretory protein. Unlike SARS-CoV-2 ORF8, SARS ORF8a could only be secreted through the conventional secretory pathway. Notably, the conventionally secreted SARS ORF8a is not glycosylated and is able to induce the release of pro-inflammatory factors. These differences might be due to the fact that ORF8 is the only SARS-CoV-2 protein with as low as approximate 20% homology to SARS(*42*). In this context, the very different secretory pattern and functional role of ORF8 in SARS-CoV-2 can partly explain why the disease spectrum of COVID-19 is different from that of SARS. Based on our data, we propose a potential competition between host and guest regarding the ORF8 protein: Primitive SARS invades host cells and secretes ORF8 through the conventional ER-Golgi trafficking pathway. ORF8 is not glycosylated in the Golgi apparatus and can induce the release of pro-inflammatory factors. However, the host does not just surrender to viral infection. During the evolutionary process, the host blocks the induction of IL-17 signaling though ORF8 glycosylation to maintain homeostasis. However, life finds a way; an evolved coronavirus develops a new pathway to bypass the glycosylation modification and secrete unglycosylated ORF8 protein, thus regaining an advantageous position in this competition for the moment. Nevertheless, this storyline needs to be validated by further studies.

In the current work, we identified YIF1B, a five-transmembrane protein, as a key molecule that mediates transport of cargo into vesicles to facilitate the unconventional secretion pattern of ORF8. Knockout of YIF1B in epithelial cells prevents the unconventional secretion of ORF8, and exogenous supplementation of YIF1B rescues the secretion of ORF8 even when Golgi apparatus is destructed by Brefeldin A. The *in vitro* translocation assay proved that YIF1B promotes the translocation of ORF8 into vesicles directly. Generally, the translocation process of leaderless cargoes into vesicles depends on chaperones(*53*). For example, leaderless cargoes bind to chaperone HSP90A that likely recognizes and unfolds the cargoes in the TMED10-channeled unconventional protein secretion pathway(*40*). Whether the translocation of leaderless ORF8 into vesicles needs chaperones in addition to the YIF1B channel is not yet known. Notably, the proteinase K protection assays shows that ORF8 cannot be translocated into vesicles without the supplementation of cell lysates containing non-inner membranes. This result does suggest the necessity of chaperones in the translocation of leaderless ORF8. It would be interesting to further investigate the specific role of chaperones in the unconventional secretion of SARS-CoV-2 ORF8.

## Materials and Methods

### Ethic statements

This study was carried out in strict accordance with the Guidelines for the Care and Use of Animals of Chongqing University. All animal experimental procedures were approved by the Animal Ethics Committees of the School of Life Sciences, Chongqing University.

### Mice

B6.Cg-Tg(K18-ACE2)2Prlmn/J mice (hACE2) were obtained from The Jackson Laboratory. To generate YIF1B-deficient (*Yif1b*^-/-^) mice based on K18-ACE2 transgenic background, we designed sgRNAs targeting the Exon 3-5 of *Yif1b*. Cas9 and sgRNAs (sgRNA1: CAGCTAACACGGTGGGTTGT, sgRNA2: GAGGCCAAGAGCCATCAGAG) were co-injected into fertilized eggs of hACE2 mice. PCR followed by sequencing analysis and western blotting were used for validation. All animal study protocols were reviewed and approved by Chongqing University School of Life Sciences review boards for animal studies.

### Cell lines and coronavirus

Calu-3 epithelial cells (HTB-55), Jurkat cells (Clone E6-1, TIB-152), THP-1 cells (TIB-202), Vero E6 cells (CRL-1586) and Sf9 cells (CRL-1711) were purchased from ATCC. HEK-293FT cells (R70007) were purchased from Thermo Fisher Scientific. Differentiation of THP-1 monocytes to macrophages was induced by 15 ng/mL phorbol 12-myristate 13-acetate (PMA) as previously described(*54*). *Ace2*^-/-^ Calu-3 epithelial cell line, *Il17ra*^-/-^ THP-1-derived macrophage cell line and YIF1B-KO Calu-3 epithelial cell line were generated using CRISPR-Cas9 system with short guide RNA sequences (*Ace2*^-/-^ cell line: ACAGTTTAGACTACAATGAG; *Il17ra*^-/-^ cell line: TGTCCATTCGATGTGAGCCA; YIF1B-KO cell line: GAGAGGCTGCAGGATAACTC). Murine alveolar epithelial cells (AECs) were isolated using the method developed by Corti and colleagues with motifications(*55, 56*). Plasmid and siRNA transfections were performed using a LONZA 4D-Nucleofector system according to the manufacturer’s instruction. The SARS-CoV-2 virus was generated by using a reverse genetic method as previously described(*25–27*). In brief, seven different DNA fragments spanning the entire genome of SARS-CoV-2 (USA_WA1/2020 SARS-CoV-2 sequence, GenBank accession No. MT020880) were synthesized by Beijing Genomics Institute (BGI, Shanghai, China) and cloned into the pUC57 or pCC1 (kindly provided by Dr. Yonghui Zheng) plasmid by standard molecular cloning methods. The sequences of the F1∼F7 fragments and the restriction enzymes used for digestion and ligation were shown in our previous study(*25*). Full-length cDNA assembly and recombinant SARS-CoV-2 virus recovery were performed as previously described(*25–27*). Virus titer was determined using a standard TCID_50_ assay. For the generation of SARS-CoV-2 ORF8-N78Q variant, AAT→CAA nucleotide substitutions were introduced into a subclone of pUC57-F7 containing the ORF8 gene of the SARS-CoV-2 wild-type infectious clone by overlap-extension PCR. Primers are as follows: F-TCAGTACATCGATATCGGT<u>CAA</u>TAT, R-GTAAACAGGAAACTGTATA<u>TTG</u>ACC.

### Viral infection

Specific-pathogen-free, ten-week-old male mice were inoculated intranasally with SARS-CoV-2 virus with 4×10^5^ plaque-forming units (PFU). Cells were infected with SARS-CoV-2 virus at a dosage of 10^5^ TCID_50_/mL within the indicated time. Cell culture supernatant obtained from infected epithelial cells was filtered by Ultipor VF Grade UDV20 Virus Removal Filter Cartridges (PALL) to remove virion, and used for macrophage stimulation. All experiments with infectious virus were performed under biosafety level 3 (BSL3+) conditions.

### Collection of secretory proteins and immunoblot

Cell culture medium was collected and centrifuged twice to remove cell debris. The supernatant was concentrated through a 10 kDa Amicon-Ultra centrifugal tube (Millipore) and prepared for immunoblot. GAPDH was used to indicate the absence of cell lysates. Immunoblot analysis was performed as previously described(*57*). Blots were probed with the indicated antibodies: anti-ORF8 (NBP3-05720), anti-GAPDH (NBP2-27103) (Novus Biologicals), anti-3CL pro (GTX135470) (GeneTex), anti-NEF (MA1-71507), anti-IL17RA (PA5-47199) (Thermo Fisher Scientific), anti-GFP (ab6556), anti-YIF1B (ab188127), anti-SEC22B (ab241585), anti-V5 (ab9137) (Abcam), anti-PDI (2446) (Cell Signaling Technology), anti-Flag (AF0036), anti-His (AF5060), anti-GST (AF5063), anti-HA (AF0039) (Beyotime Biotechnology).

### Enzyme linked immunosorbent assay

Cell culture supernatant was purified by centrifugation, and assayed by enzyme linked immunosorbent assays (ELISA). ORF8 antibody (NBP3-05720, Novus Biologicals) was coated with blank ELISA plates in carbonate buffer to prepare ELISA kit.

Cytokine and chemokine ELISA kits were purchased from eBiosciences (Thermo Fisher Scientific).

### Block of conventional secretory pathway

Brefeldin A (00-4506-51, eBioscience, 3 μg/mL) or Monensin (00-4505-51, eBioscience, 2 μM) was added in the culture medium of epithelial cells for 2 hours to inhibit the ER-Golgi trafficking and block the ER-Golgi conventional secretion pathway.

### N-linked glycosylation identification by HPLC-MS/MS

N-linked glycosylation identification of SARS-CoV-2 ORF8 protein was performed as previously described with slight modifications(*58*). Firstly, Calu-3 epithelial cells were infected by SARS-CoV-2 with/without Brefeldin A or Monensin pretreatment. The supernatant was collected and concentrated through a 10 kDa Amicon-Ultra centrifugal tube (Millipore). Then, the N-glycopeptides were enriched with Zic-HILIC (Fresh Bioscience), eluted and dried for deglycosylation. Enriched N-glycopeptides were digested using PNGase F dissolved in 50 mM NH_4_HCO_3_ for 2 hours at 37 °C to remove N-linked glycosylation. Finally, deglycosylated peptides were dissolved in 0.1% FA for tandem mass spectrum analysis. MS1 was analyzed at an Orbitrap resolution of 120,000 using a scan range (m/z) of 800 to 2000 (N-glycopeptides before and after enrichment), or 350 to 1550 (deglycosylated peptides). The RF lens, AGC target, maximum injection time, and exclusion duration were 30%, 2.0 e^4^, 100 ms and 15 s, respectively. MS2 was analyzed with an isolation window (m/z) of 2 at an Orbitrap resolution of 15,000. The AGC target, maximum injection time, and the HCD type were standard, 250 ms, and 30%, respectively.

### Deglycosylation assays

Tunicamycin (12819, Cell Signaling Technology, 2µg/mL) was added into epithelial cells to prevent N-linked glycosylation. PNGase F (P0704, NEB, 1,000 units/µg protein) was used to remove the N-linked glycosylation in purified ORF8 protein. 1 µg glycoprotein, 2 µL GlycoBuffer 2 (10×), 2 µL PNGase F and H_2_O were mixed to form a 20 µL mixture. The mixture was incubated at 37°C for 8 hours, followed by western blotting analysis. To remove O-linked glycosylation, purified proteins were directly digested by O-glycosidase (P0733, NEB, 4,000 units/µg protein) and α2-3, 6, 8, 9 Neuraminidase A (P0722, NEB, 4 units/µg protein). 10 µg glycoprotein, 1 µL 10× Glycoprotein Denaturing Buffer and H_2_O were combined to make a 10 µL mixture. The glycoprotein was denatured by heating the mixture at 100°C for 10 min. Then, 2 µL 10× GlycoBuffer 2, 2 µL 10% NP40, 2 µL α2-3, 6, 8, 9 Neuraminidase A, 1 µL O-Glycosidase and H_2_O were mixed to form a total volume of 20 µL and incubated at 37°C for 4 hours, followed by western blotting analysis.

### Peptide synthesis and artificial glycosylation modification

SARS-CoV-2 ORF8 peptide was synthesized according to the NCBI published sequence (accession number: YP_009724396.1). Fmoc-L-Asn ((Ac)3-β-D-GlcNAc)-OH modification was performed by Shanghai Science Peptide Biological Technology Co., Ltd. ORF8 peptides with or without N-glycosylation were dissolved in 40 µL DMSO and diluted with PBS buffer for cell stimulation (1 µg/mL) or mice aerosol infection (10 µg/g).

### Identification of the interaction channel proteins of ORF8

DHFR assay followed by HPLC-MS/MS was used to screen the channel proteins mediating ORF8 translocation in an unconventional secretion pathway as previously described(*12*). Calu-3 cells expressing ORF8-Flag-DHFR were treated with Brefeldin A (3 μg/mL) or Monensin (2 μM) to inhibit conventional secretion pathways. Aminopterin (15 μM) was added to inhibit DHFR unfolding. After crosslinking, membrane fractions were collected, lysed and immunoprecipitated with anti-Flag beads for HPLC-MS/MS analysis.

### Duolink PLA assay

Duolink PLA assay was performed according to the kit manual (Duolink In Situ Detection Reagents Red, DUO92008, Sigma). In brief, Flag-tagged ORF8 and HA-tagged YIF1B were co-transfected into HEK-293FT cells. After 12 hours, cells were fixed with 4% paraformaldehyde (PFA) for 20 min and permeabilized with 0.2% Triton X-100. Next, the fixed cells were blocked, incubated with antibodies and PLA probes according to the protocol. Images were captured followed by quantification using Image J software.

### *In vitro* translocation assay

Purified YIF1B proteins (NCBI accession number: NP_001034761.1) was synthesized by MembraneMax^TM^ cell-free protein synthesis system according to the manufacturers’ protocol (A10633, Thermo Fisher Scientific). Purified ORF8 protein was expressed in baculovirus expression vector system (Thermo Fisher Scientific). In brief, ORF8 protein-coding sequence without signal peptide was optimized according to the insect cell codon preference, and synthesized by Sangon Biotech. The Asparagine 78 was mutated to Glutarnine to prevent glycosylation. Then, synthesized ORF8 sequence was amplified by PCR and inserted into a pFastBac HT A plasmid (Thermo Fisher Scientific). Recombinant pFastBac-ORF8 plasmid was transformed into DH10Bac-competent cells to obtain Bacmid-ORF8, which was transfected into Sf9 cells to produce ORF8 protein. Total lipids were extracted from HEK-293FT cells as previously described(*12*). Purified YIF1B and lipids were reconstituted into proteolioposomes. To be specific, lipids were repeatedly frozen and thawed in a 42°C water bath for 10 times. Triton X-100 was used to dilute lipids to a final concentration of 0.05%, and the lipid solution was rotated at 4°C for 30 min. Then, the purified YIF1B protein was added into the lipid solution and rotated for 1 hour, followed by incubation with Biobeads SM2 (Bio-Rad Laboratories) to absorb the detergent. After centrifugation to remove the beads, a membrane flotation procedure was performed to collect top fractions containing proteolioposomes. Next, proteolioposomes were mixed with ORF8 protein in HEK-293FT lysates (whole cell lysates were centrifuged at 1,000×g for 10 min and the supernatant was ultra-centrifuged at 100,000×g for 40 min to remove endomembrane) for 1 hour. The mixture was centrifuged, aliquoted into three parts, and collected for proteinase K protection test.

### Proteinase K protection test

Proteinase K protection test was performed as previously described(*40*). Briefly, cells were harvested and lysed in HB1 buffer (20 mM HEPES-KOH, pH 7.2, 400 mM sucrose, and 1 mM EDTA) with 0.3 mM DTT and protease inhibitors. After centrifugation, supernatant was ultra-centrifuged at 100,000×g for 40 min to collect the total membrane pellet, followed by membrane flotation assay. The membrane fraction floating on the top was collected and divided into three parts (without proteinase K, with proteinase K, or with proteinase K and 0.5% Triton X-100). The reactions were performed on ice and stopped by adding PMSF and SDS loading buffer. The samples were immediately heated at 98°C for 5 min, followed by SDS-PAGE. Protein disulfide isomerase (PDI) and a vesicle trafficking protein, SEC22 Homolog B (SEC22B), were used as the positive and negative control, respectively.

### H&E staining

Mice were anaesthetized with isoflurane, and lung lobes were harvested at indicated time points. Tissues were fixed with 10% PFA for more than 24 hours and embedded in paraffin. The paraffin blocks were cut into 2 μm-thick sections and stained using a standard Hematoxylin and eosin (H&E) procedure.

### Statistical Analysis

Sample size was based on empirical data from pilot experiments. The investigators were blinded during data collection and analysis. A value of P < 0.05 was considered significant.

## Acknowledgments

This work was supported by the National Natural Science Foundation of China, SGC’s Rapid Response Funding for COVID-19 (C-0002), National Natural Science Foundation of China (No. 81970008, 82000020, 31702205), the Fundamental Research Funds for the Central Universities (No. 2021CDJZYJH-002, 2019CDYGZD009 and 2020CDJYGRH-1005), Natural Science Foundation of Chongqing, China (cstc2020jcyj-msxmX0460 and cstc2020jcyj-bshX0105) and Chongqing Talents: Exceptional Young Talents Project (No. cstc2021ycjh-bgzxm0099). The funders had no role in study design, data collection and analysis, decision to publish, or preparation of the manuscript.

## Author contributions

H. Wu, X. Lin and B. Fu conceived and designed the study. H. Wu, X, Lin, B. Fu, Y. Xiong, N. Xing, W. Xue, D. Guo, M. Y. Zaky, K. C. Pavani, D. Kunec and J. Trimpert performed the experiments. H. Wu, X. Lin and B. Fu analyzed the data. H. Wu, X. Lin, B. Fu and J. Trimpert wrote the manuscript. All authors read and approved the final manuscript.

## Figure legends

**Figure S1 SARS-CoV-2 ORF8 can be secreted without the presence of complete virus**

(A) Validation of Calu-3 *Ace2*^+/+^, Calu-3 *Ace2*^-/-^, THP-1 DM *Il17ra*^+/+^ and THP-1 DM *Il17ra*^-/-^ cells. The expressions of ACE2 and IL17RA were detected by western blotting. Representative images from n = 3 biological replicates are shown.

(B) Schematic diagram of THP-1 DM cells stimulation model. HEK-293FT cells were transfected with Flag-tagged ORF8, 3CL pro, or Nef. After 12 hours, the supernatant was collected and divided into two parts. One part was used to purify secretory proteins, followed by western blotting; the other part was used to stimulate THP-1 DM cells for 12 hours. The release of cytokines and chemokines was detected by ELISA.

(C) Secretory proteins obtained from HEK-293FT cell supernatant in (B) were detected by western blotting. Representative images from n = 3 biological replicates are shown.

(D) The release of cytokines and chemokines from THP-1 DM cells in (B) was detected by ELISA. Data are shown as the mean ± s.e.m. of n = 3 biological replicates.

One-way ANOVA followed by Bonferroni post *hoc* test (D)was used for data analysis. *, p < 0.05, **, p < 0.01.

**Figure S2 SARS ORF8a cannot be glycosylated**

(A, B) Brefeldin A (3 μg/mL) or Monensin (2 μM) was used to pretreat Calu-3 cells for 2 hours, followed by SARS-CoV-2 infection for 12 hours at a dosage of 10^5^ TCID_50_/mL (A), or Flag-tagged ORF8 plasmid transfection for 12 hours (B). The supernatant was collected to stimulate THP-1 DM cells within the specified time. The release of cytokines and chemokines was detected by ELISA. Data are shown as the mean ± s.e.m. of n = 3 biological replicates.

(C) PNGase F (1,000 units/µg protein), O-Glycosidase (4,000 units/µg protein) or α2-3, 6, 8, 9 Neuraminidase A (4 units/μg glycoprotein) was added into purified SARS ORF8a protein to release glycans. Western blotting was used to detect the glycosylation of ORF8a protein. Representative images from n = 3 biological replicates are shown.

(D) Calu-3 cells were transfected with SARS ORF8a-Flag plasmids. Tunicamycin (2 µg/mL) was added into Calu-3 cells for 2 hours to prevent N-linked glycosylation, or PNGase F (1,000 units/µg protein) was used to remove the N-linked glycosylation in purified ORF8 protein. After deglycosylation assays, the cell culture supernatant or purified ORF8a protein was used to stimulate THP-1 DM cells. After 12 hours, the release of cytokines and chemokines was detected by ELISA. Data are shown as the mean ± s.e.m. of n = 3 biological replicates.

Two-way (A, B) or one-way (D) ANOVA followed by Bonferroni post *hoc* test was used for data analysis. Abbreviations: n.s., not significant.

**Figure S3 ORF8 N78 glycosylation blocks the activation of IL-17 pathway**

(A) Calu-3 cells were infected with SARS-CoV-2 ORF8-N78Q variant, or transfected with ORF8-N78Q plasmids. Twelve hours later, the supernatant was used to stimulate THP-1 DM cells for another 12 hours. The activation of IL-17 pathway was evaluated by testing NF-κB activity. Data are shown as the mean ± s.e.m. of n = 3 biological replicates.

(B, C) Calu-3 cells were infected with SARS-CoV-2 ORF8-N78Q variant (B), or transfected with ORF8-N78Q plasmids (C). The supernatant was collected to purify ORF8 protein. After PNGase F digestion, the ORF8 protein was used to stimulate THP-1 DM cells. After 12 hours, the activation of IL-17 pathway was evaluated by testing NF-κB activity. Data are shown as the mean ± s.e.m. of n = 3 biological replicates.

(D, E) Brefeldin A (3 μg/mL) or Monensin (2 μM) was used to pretreat Calu-3 cells for 2 hours, followed by infection with SARS-CoV-2 ORF8-N78Q variant (D) or transfection with ORF8-N78Q plasmids (E) for 12 hours. The supernatant was used to stimulate THP-1 DM cells for 12 hours. The secretion of ORF8 was detected by ELISA. Data are shown as the mean ± s.e.m. of n = 3 biological replicates.

(F, G) The activation of IL-17 pathway in (D, E) was evaluated by testing NF-κB activity. Data are shown as the mean ± s.e.m. of n = 3 biological replicates. Two-way ANOVA followed by Bonferroni post *hoc* test was used for data analysis. *, p < 0.05, **, p < 0.01. Abbreviations: n.s., not significant.

**Figure S4 SARS-CoV-2 ORF8 can be secreted through an unconventional pathway**

(A) Calu-3 cells were infected with SARS-CoV-2 for 12 hours at a dosage of 10^5^ TCID_50_/mL. Starvation assay (12 hours) or Rapamycin (50 nM) was used to induce autophagy. The secretion of ORF8 was detected by western blotting. Brefeldin A (3 μg/mL) was used to block conventional secretion of ORF8. Representative images from n = 3 biological replicates are shown.

(B) Calu-3 cells were infected with SARS-CoV-2 for 12 hours at a dosage of 10^5^ TCID_50_/mL. 3-MA (10 mM) or Wtm (20 nM) was used to inhibit the autophagy induced by starvation. The secretion of ORF8 was detected by western blotting. Brefeldin A (3 μg/mL) was used to block conventional secretion of ORF8. Representative images from n = 3 biological replicates are shown.

(C) Calu-3 cells were infected with SARS-CoV-2 for 12 hours at a dosage of 10^5^ TCID_50_/mL. siRNA of *Atg5*, *Atg2a*, or *Atg2b* was transfected into cells to inhibit the autophagy induced by starvation. The secretion of ORF8 was detected by western blotting. Brefeldin A (3 μg/mL) was used to block conventional secretion of ORF8. Representative images from n = 3 biological replicates are shown.

(D) Calu-3 cells were infected with SARS-CoV-2, or transfected with ORF8-Flag. After 12 hours, the co-localization of ORF8 and ERGIC-53 was observed by immunofluorescence. Brefeldin A (3 μg/mL) was used to block conventional secretion of ORF8. Representative images from n = 3 biological replicates are shown. Scale bar = 10 μm.

(E) Calu-3 cells were infected with SARS-CoV-2 for 12 hours at a dosage of 10^5^ TCID_50_/mL. Whole cell lysates were collected for Proteinase K protection assay. PDI and SEC22B were used as positive control and negative control, respectively. Brefeldin A (3 μg/mL) was used to block conventional secretion of ORF8. Representative images from n = 3 biological replicates are shown.

(F) Schematic diagram of screening channel proteins secreted by ORF8 through an unconventional pattern. Brefeldin A (3 μg/mL) or Monensin (2 μM) was used to pretreat HEK-293FT cells for 2 hours, followed by transfection with DHFR/Flag-tagged ORF8 for 12 hours. Aminopterin was used to inhibit DHFR unfolding. Whole cell lysates were collected for HPLC-MS/MS analysis. Transmembrane proteins interacting with ORF8 both in Monensin and Brefeldin A treatment groups were considered as candidates.

(G) Calu-3 cells were infected with SARS-CoV-2 for 12 hours at a dosage of 10^5^ TCID_50_/mL. siRNAs of potential channel proteins were transfected into cells, and the secretion of ORF8 was detected by western blotting. Representative images from n = 3 biological replicates are shown.

**Figure S5 Validation of *Yif1b*^-/-^ mice**

(A) Schematic diagram of knockout strategy in generation of *Yif1b*^-/-^ hACE2 mice using CRISPR-Cas9. Two sgRNAs were designed to delete 3-5 exons of *Yif1b*.

(B) PCR validation of *Yif1b*^-/-^ mice genotype. Representative images from n = 3 biological replicates are shown.

(C) The expressions of YIF1B in lungs obtained from *Yif1b*^+/+^ and *Yif1b*^-/-^ mice were detected by western blotting. Representative images from n = 3 biological replicates are shown.

## References

1. L. Yang et al., The signal pathways and treatment of cytokine storm in COVID-19. Signal Transduct Target Ther 6, 255 (2021).

2. C. Huang et al., Clinical features of patients infected with 2019 novel coronavirus in Wuhan, China. Lancet 395, 497–506 (2020).

3. X. Tang et al., On the origin and continuing evolution of SARS-CoV-2. Natl Sci Rev 7, 1012–1023 (2020).

4. D. Muth et al., Attenuation of replication by a 29 nucleotide deletion in SARS-coronavirus acquired during the early stages of human-to-human transmission. Sci Rep 8, 15177 (2018).

5. B. E. Young et al., Effects of a major deletion in the SARS-CoV-2 genome on the severity of infection and the inflammatory response: an observational cohort study. Lancet 396, 603–611 (2020).

6. X. Lin et al., ORF8 contributes to cytokine storm during SARS-CoV-2 infection by activating IL-17 pathway. iScience 24, 102293 (2021).

7. P. Bhadra, V. Helms, Molecular Modeling of Signal Peptide Recognition by Eukaryotic Sec Complexes. Int J Mol Sci 22, (2021).

8. J. McCaughey, D. J. Stephens, ER-to-Golgi Transport: A Sizeable Problem. Trends in cell biology 29, 940–953 (2019).

9. R. Z. Murray, J. L. Stow, Cytokine Secretion in Macrophages: SNAREs, Rabs, and Membrane Trafficking. Front Immunol 5, 538 (2014).

10. V. Malhotra, Unconventional protein secretion: an evolving mechanism. The EMBO journal 32, 1660–1664 (2013).

11. J. P. Steringer, W. Nickel, A direct gateway into the extracellular space: Unconventional secretion of FGF2 through self-sustained plasma membrane pores. Seminars in cell & developmental biology 83, 3–7 (2018).

12. M. Zhang et al., A Translocation Pathway for Vesicle-Mediated Unconventional Protein Secretion. Cell 181, 637–652 e615 (2020).

13. R. P. McNamara et al., Nef Secretion into Extracellular Vesicles or Exosomes Is Conserved across Human and Simian Immunodeficiency Viruses. mBio 9, (2018).

14. N. Mukhamedova et al., Exosomes containing HIV protein Nef reorganize lipid rafts potentiating inflammatory response in bystander cells. PLoS Pathog 15, e1007907 (2019).

15. M. Z. Mehboob, M. Lang, Structure, function, and pathology of protein O-glucosyltransferases. Cell Death Dis 12, 71 (2021).

16. K. W. Moremen, M. Tiemeyer, A. V. Nairn, Vertebrate protein glycosylation: diversity, synthesis and function. Nat Rev Mol Cell Biol 13, 448–462 (2012).

17. K. T. Schjoldager, Y. Narimatsu, H. J. Joshi, H. Clausen, Global view of human protein glycosylation pathways and functions. Nat Rev Mol Cell Biol 21, 729–749 (2020).

18. W. Chen, Y. Zhong, Y. Qin, S. Sun, Z. Li, The evolutionary pattern of glycosylation sites in influenza virus (H5N1) hemagglutinin and neuraminidase. PLoS One 7, e49224 (2012).

19. S. S. L. Yap, T. Nguyen-Khuong, P. M. Rudd, S. Alonso, Dengue Virus Glycosylation: What Do We Know? Front Microbiol 8, 1415 (2017).

20. A. J. Behrens, M. Crispin, Structural principles controlling HIV envelope glycosylation. Curr Opin Struct Biol 44, 125–133 (2017).

21. D. L. Carbaugh, H. M. Lazear, Flavivirus Envelope Protein Glycosylation: Impacts on Viral Infection and Pathogenesis. Journal of virology 94, (2020).

22. W. Tian et al., O-glycosylation pattern of the SARS-CoV-2 spike protein reveals an “O-Follow-N” rule. Cell research 31, 1123–1125 (2021).

23. Q. Li et al., The Impact of Mutations in SARS-CoV-2 Spike on Viral Infectivity and Antigenicity. Cell 182, 1284–1294 e1289 (2020).

24. Y. Watanabe, J. D. Allen, D. Wrapp, J. S. McLellan, M. Crispin, Site-specific glycan analysis of the SARS-CoV-2 spike. Science 369, 330–333 (2020).

25. H. Wu et al., Nucleocapsid mutations R203K/G204R increase the infectivity, fitness, and virulence of SARS-CoV-2. Cell Host Microbe, (2021).

26. J. A. Plante et al., Spike mutation D614G alters SARS-CoV-2 fitness. Nature 592, 116–121 (2021).

27. X. Xie et al., An Infectious cDNA Clone of SARS-CoV-2. Cell Host Microbe 27, 841–848 e843 (2020).

28. Z. Ke et al., Structures and distributions of SARS-CoV-2 spike proteins on intact virions. Nature 588, 498–502 (2020).

29. Q. Wang et al., Structural and Functional Basis of SARS-CoV-2 Entry by Using Human ACE2. Cell 181, 894–904 e899 (2020).

30. S. Lin, S. Pandruvada, H. Yu, Inhibition of Sphingosine-1-Phosphate Receptor 2 by JTE013 Promoted Osteogenesis by Increasing Vesicle Trafficking, Wnt/Ca(2+), and BMP/Smad Signaling. Int J Mol Sci 22, (2021).

31. E. R. McGlone et al., Receptor Activity-Modifying Protein 2 (RAMP2) alters glucagon receptor trafficking in hepatocytes with functional effects on receptor signalling. Mol Metab 53, 101296 (2021).

32. X. Zhang, Y. Wang, Nonredundant Roles of GRASP55 and GRASP65 in the Golgi Apparatus and Beyond. Trends in biochemical sciences 45, 1065–1079 (2020).

33. Y. Huang et al., FUT8-mediated aberrant N-glycosylation of B7H3 suppresses the immune response in triple-negative breast cancer. Nature communications 12, 2672 (2021).

34. C. E. Martin et al., Posttranslational modifications of serine protease TMPRSS13 regulate zymogen activation, proteolytic activity, and cell surface localization. The Journal of biological chemistry 297, 101227 (2021).

35. X. Wang et al., ER stress promotes HBV production by enhancing utilization of the autophagosome-multivesicular body axis. Hepatology, (2021).

36. M. Akbar et al., Translational targeting of inflammation and fibrosis in frozen shoulder: Molecular dissection of the T cell/IL-17A axis. Proc Natl Acad Sci U S A 118, (2021).

37. W. Nickel, C. Rabouille, Mechanisms of regulated unconventional protein secretion. Nat Rev Mol Cell Biol 10, 148–155 (2009).

38. C. Rabouille, Pathways of Unconventional Protein Secretion. Trends in cell biology 27, 230–240 (2017).

39. L. Liu, M. Zhang, L. Ge, Protein translocation into the ERGIC: an upstream event of secretory autophagy. Autophagy 16, 1358–1360 (2020).

40. M. Zhang, S. J. Kenny, L. Ge, K. Xu, R. Schekman, Translocation of interleukin-1beta into a vesicle intermediate in autophagy-mediated secretion. Elife 4, (2015).

41. S. Subramani, V. Malhotra, Non-autophagic roles of autophagy-related proteins. EMBO Rep 14, 143–151 (2013).

42. T. G. Flower et al., Structure of SARS-CoV-2 ORF8, a rapidly evolving immune evasion protein. Proc Natl Acad Sci U S A 118, (2021).

43. D. Kamiyama et al., Versatile protein tagging in cells with split fluorescent protein. Nature communications 7, 11046 (2016).

44. L. Zhang et al., Crystal structure of SARS-CoV-2 main protease provides a basis for design of improved alpha-ketoamide inhibitors. Science 368, 409–412 (2020).

45. D. Chen et al., ORF3a of SARS-CoV-2 promotes lysosomal exocytosis-mediated viral egress. Dev Cell, (2021).

46. Y. Ren et al., The ORF3a protein of SARS-CoV-2 induces apoptosis in cells. Cellular & molecular immunology 17, 881–883 (2020).

47. I. Kimura et al., Sarbecovirus ORF6 proteins hamper induction of interferon signaling. Cell Rep 34, 108916 (2021).

48. A. Hachim et al., ORF8 and ORF3b antibodies are accurate serological markers of early and late SARS-CoV-2 infection. Nat Immunol 21, 1293–1301 (2020).

49. A. Grifoni et al., Targets of T Cell Responses to SARS-CoV-2 Coronavirus in Humans with COVID-19 Disease and Unexposed Individuals. Cell 181, 1489–1501 e1415 (2020).

50. Y. Zhang et al., The ORF8 protein of SARS-CoV-2 mediates immune evasion through down-regulating MHC-Iota. Proc Natl Acad Sci U S A 118, (2021).

51. K. K. To et al., Temporal profiles of viral load in posterior oropharyngeal saliva samples and serum antibody responses during infection by SARS-CoV-2: an observational cohort study. Lancet Infect Dis 20, 565–574 (2020).

52. F. X. Lescure et al., Clinical and virological data of the first cases of COVID-19 in Europe: a case series. Lancet Infect Dis 20, 697–706 (2020).

53. S. Kaushik, A. M. Cuervo, Chaperone-mediated autophagy: a unique way to enter the lysosome world. Trends in cell biology 22, 407–417 (2012).

54. I. Pantazi et al., SARS-CoV-2/ACE2 Interaction Suppresses IRAK-M Expression and Promotes Pro-Inflammatory Cytokine Production in Macrophages. Front Immunol 12, 683800 (2021).

55. M. Corti, A. R. Brody, J. H. Harrison, Isolation and primary culture of murine alveolar type II cells. Am J Respir Cell Mol Biol 14, 309–315 (1996).

56. L. Cakarova et al., Macrophage tumor necrosis factor-alpha induces epithelial expression of granulocyte-macrophage colony-stimulating factor: impact on alveolar epithelial repair. Am J Respir Crit Care Med 180, 521–532 (2009).

57. B. Fu et al., MiR-342 controls Mycobacterium tuberculosis susceptibility by modulating inflammation and cell death. EMBO Rep 22, e52252 (2021).

58. Y. Zhang et al., Site-specific N-glycosylation Characterization of Recombinant SARS-CoV-2 Spike Proteins. Mol Cell Proteomics, 100058 (2020).

