## Supplementary figures and images for "Unconventional secretion of unglycosylated ORF8 is critical for the cytokine storm during SARS-CoV-2 infection"

### Supplemental Figure 1

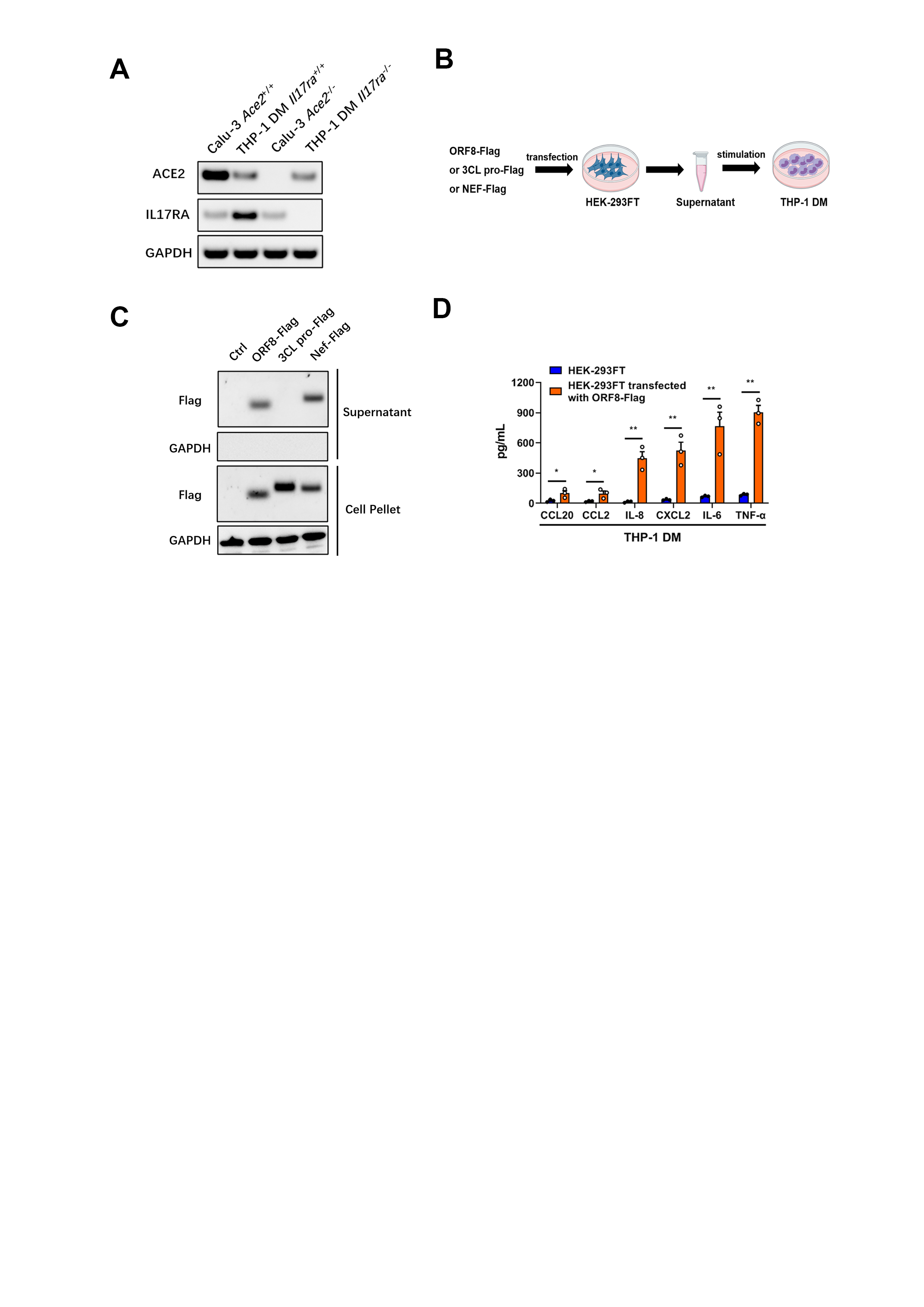

### Supplemental Figure 2

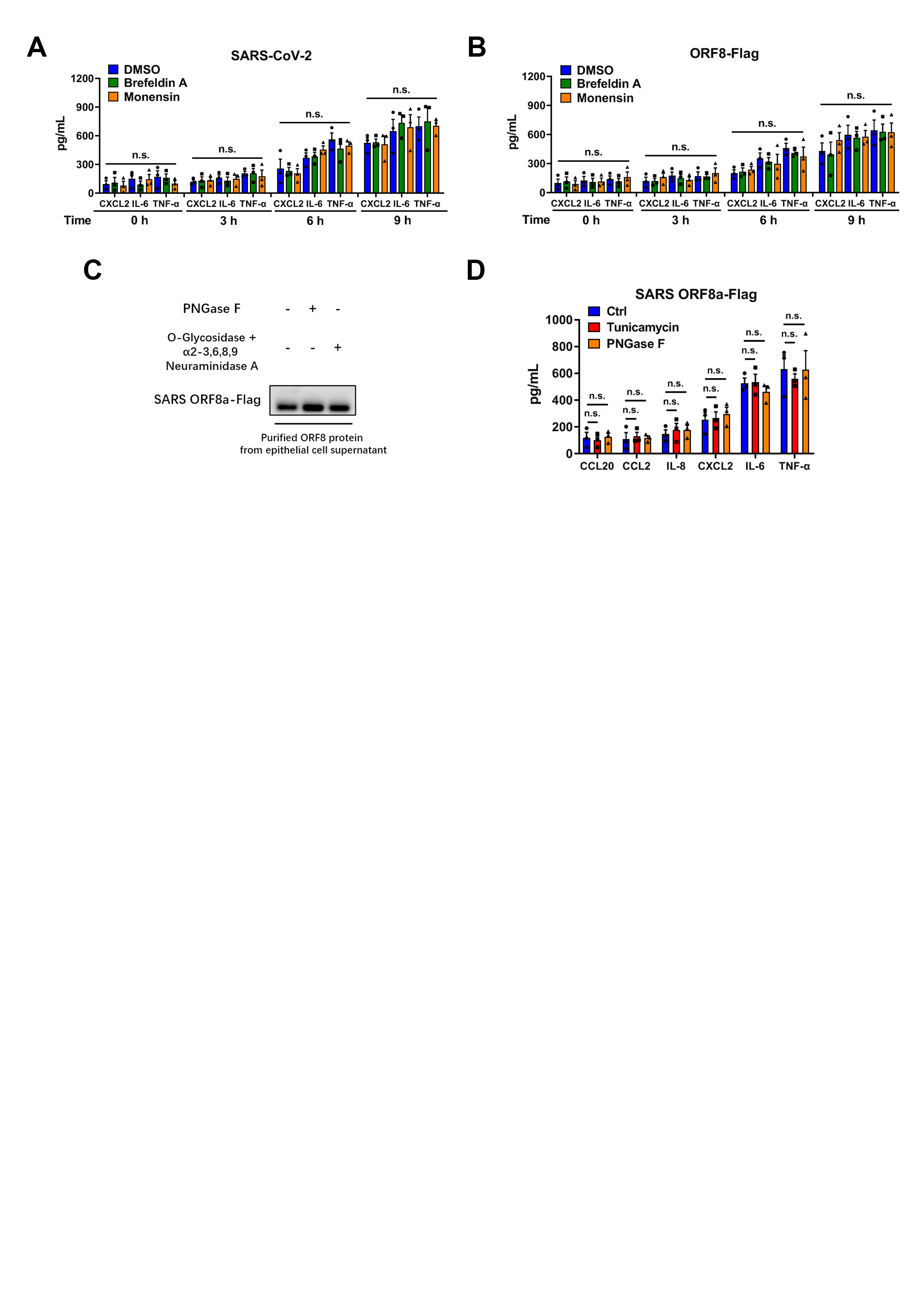

### Supplemental Figure 3

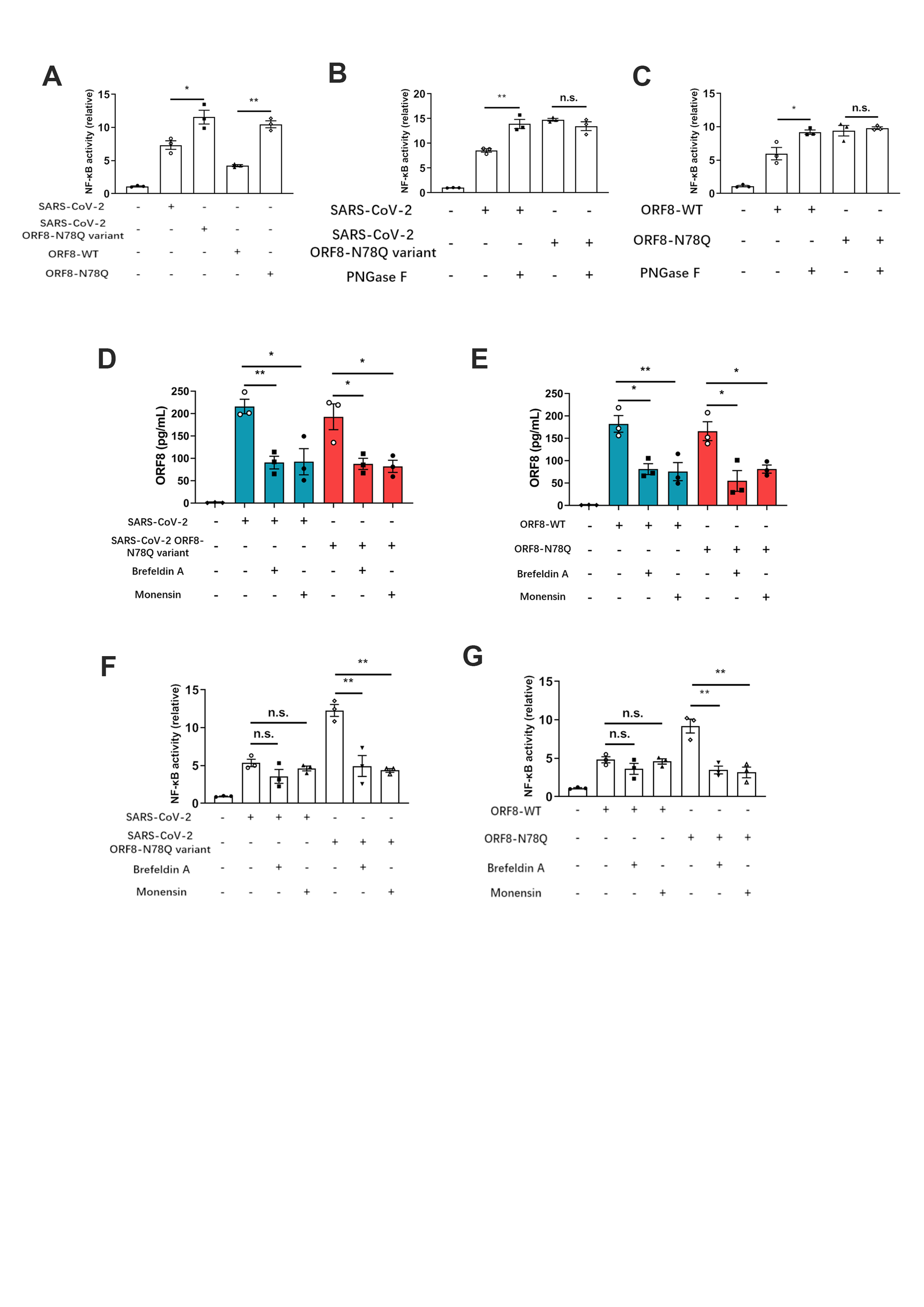

### Supplemental Figure 4

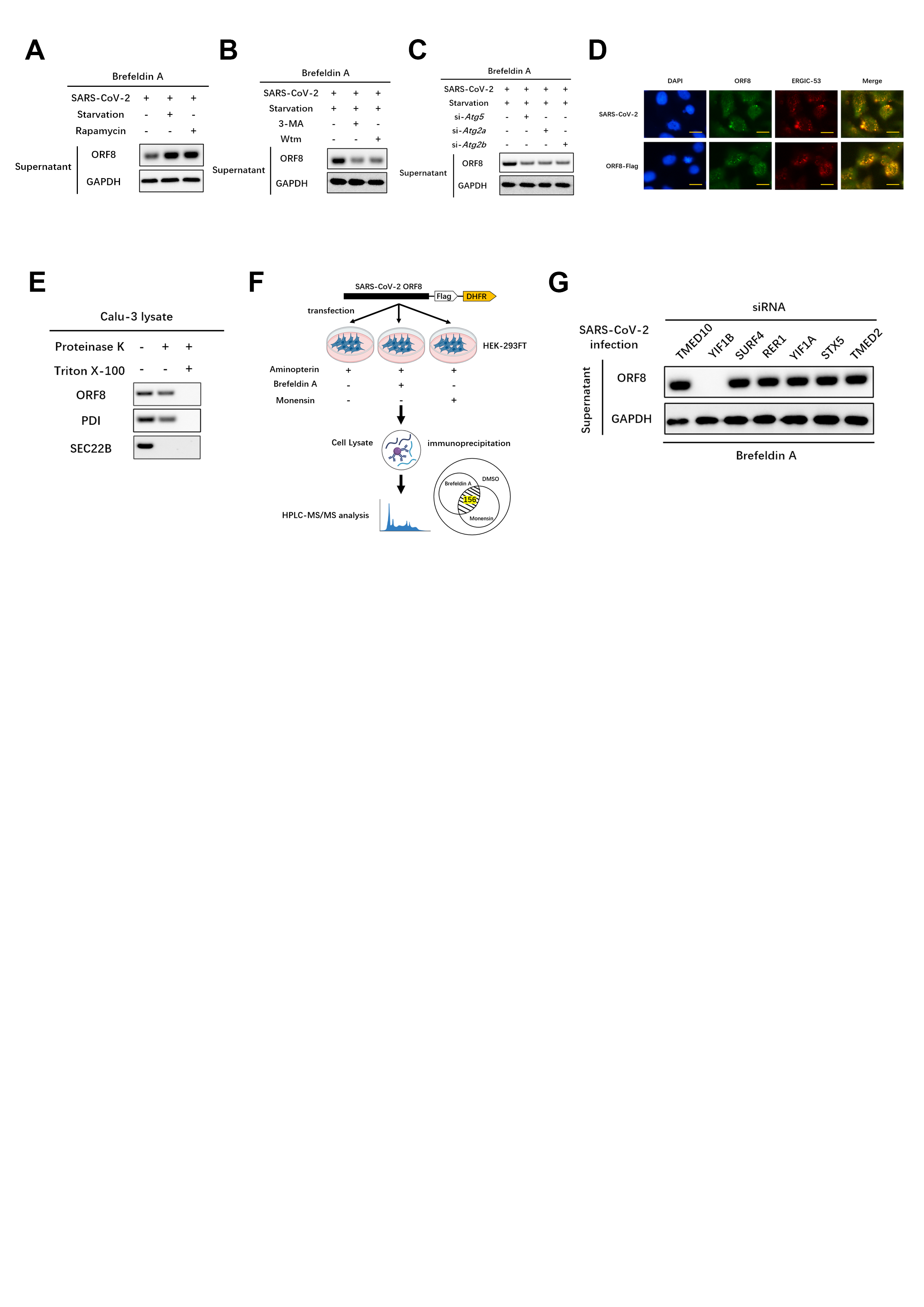

### Supplemental Figure 5

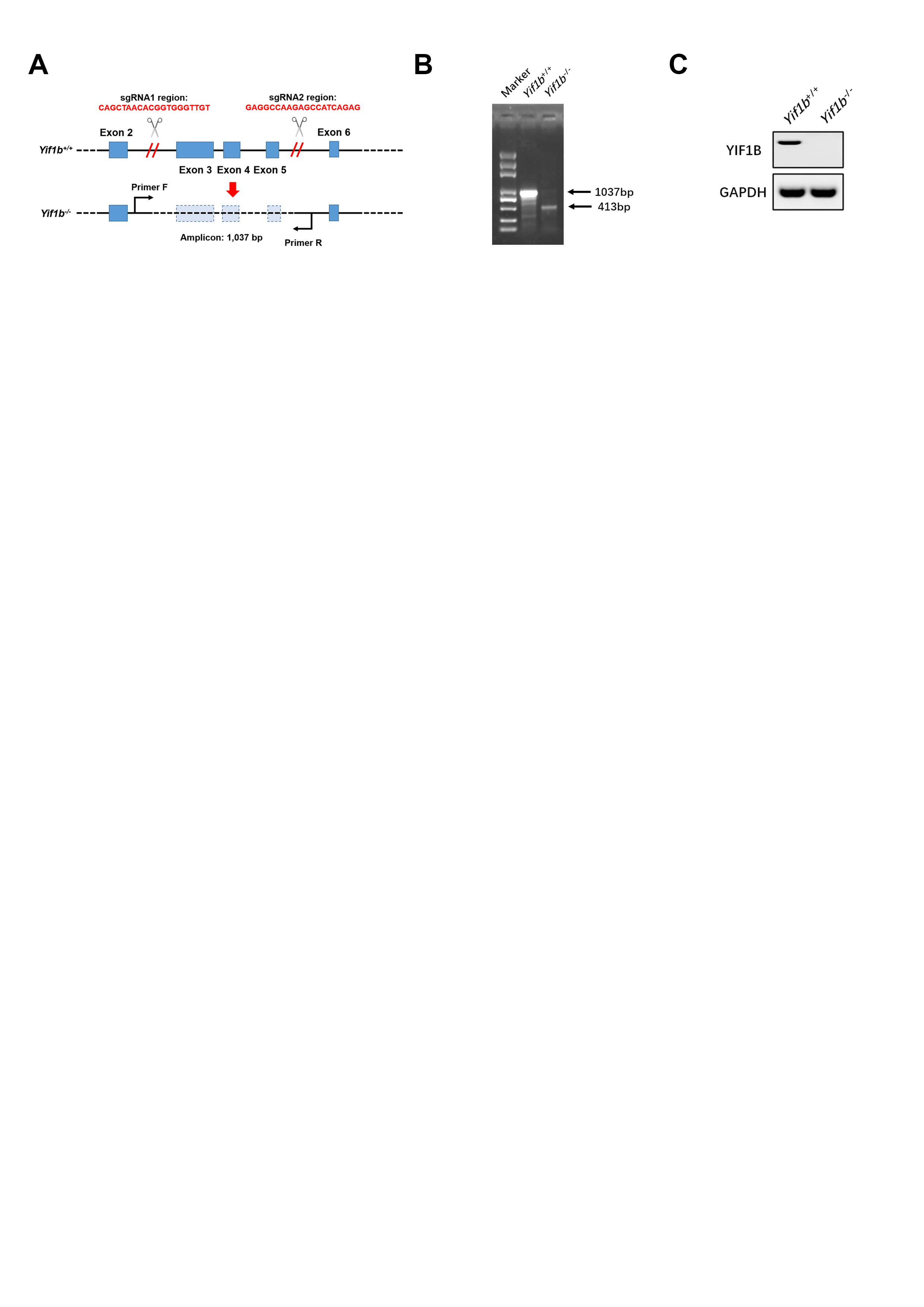
